# Individual differences in the effects of musical familiarity and musical features on brain activity during relaxation

**DOI:** 10.1101/2024.11.27.625650

**Authors:** Rebecca Jane Scarratt, Martin Dietz, Peter Vuust, Boris Kleber, Kira Vibe Jespersen

## Abstract

Finding a way to relax is increasingly difficult in our overstimulating modern society and chronic stress can have severe psychological and physiological consequences. Music is a promising tool to promote relaxation by lowering heart rate, modulating mood and thoughts, and providing a sense of safety. Here, we used functional magnetic resonance imaging (fMRI) to investigate how music influence brain activity during relaxation with a particular focus on the participants’ experience of different types of music. In a 2×2 design, 57 participants were scanned while rating how relaxed they felt after listening to 28-second excerpts of either familiar or unfamiliar relaxation music with calm or energetic features. Behaviourally, calm music was the strongest predictor of relaxation, followed by familiar music. fMRI results revealed activations of auditory, motor, emotion, and memory areas, for listening to familiar compared to unfamiliar music. This suggests increased audio-motor synchronization and participant engagement of known music. Listening to unfamiliar music was correlated with attention-related brain activity, suggesting increased attentional load for this music. Behaviourally, we identified four clusters of participants based on their relaxation response to the different types of music. These groups also displayed distinct auditory and motor activity patterns, suggesting that the behavioural responses are linked to changes in music processing. Interestingly, some individuals found energetic music to be relaxing if it is familiar, whereas others only found calm music to be relaxing. Such individual behavioural and neurological differences in relaxation responses to music emphasise the importance of developing personalised music-based interventions.

**Graphical Abstract:** 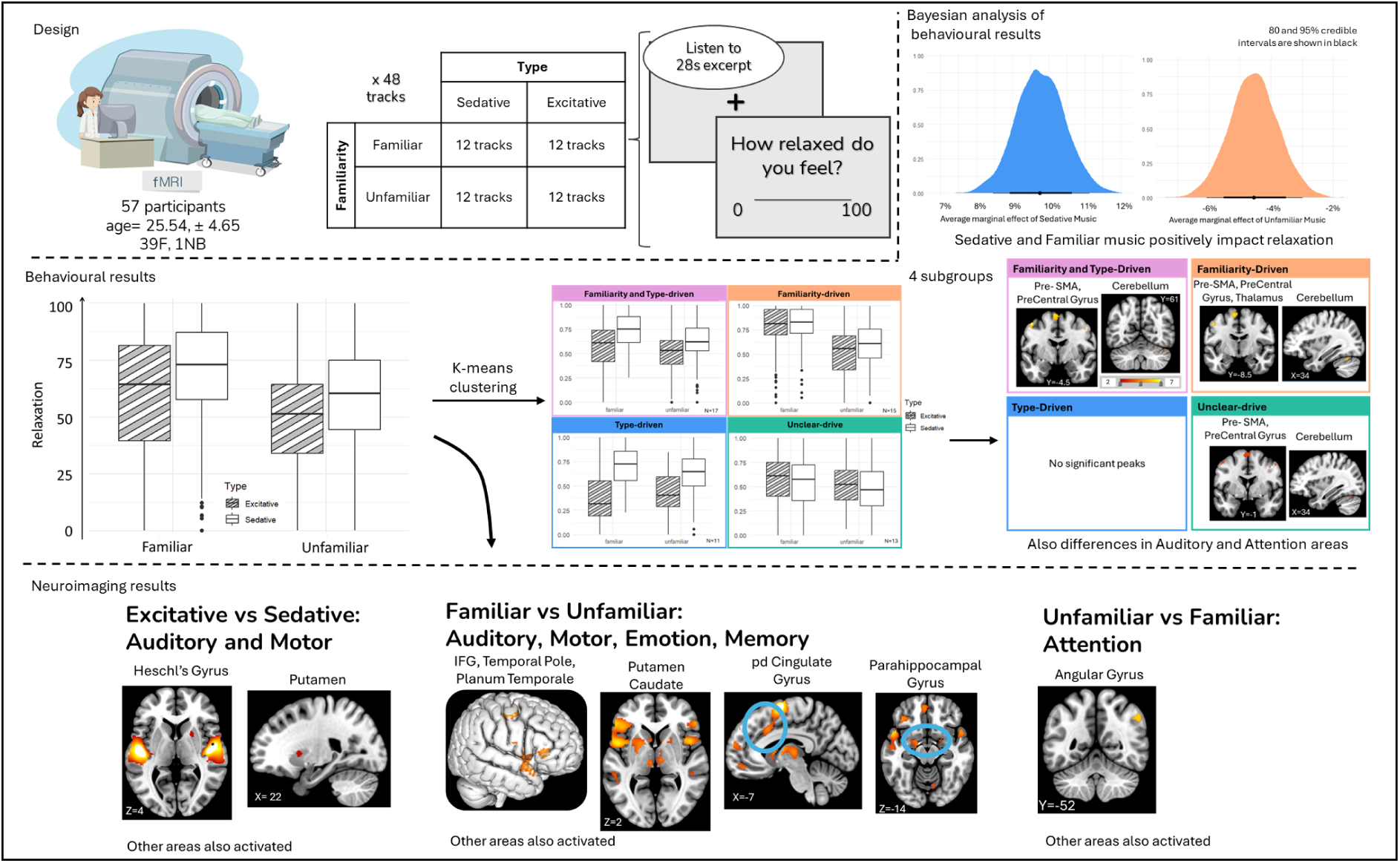

**Highlights:**

- Participants listen to relaxation tracks varying in familiarity and energy levels
- Sedative music followed by familiar music increased relaxation ratings the most
- Auditory, motor, memory and attention activations depending on the music type
- 4 distinct clusters of behavioural responses to familiar or sedative music
- These groups are also present in brain activity

## Introduction

Music is used by many to induce relaxation and relieve stress, in both adults and children in a variety of different situations (Aitken et al., 2002; Esch et al., 2003; Lai et al., 2008; Lesiuk, 2008; Nilsson, 2008; Sung & Chang, 2005; Tan et al., 2012). When subjected to stress, the body engages the stress response, a state of hyper arousal with increase heart rate, respiration rate, and involvement of the sympathetic nervous system: the fight-flight response (Esch et al., 2003). To go back to baseline state, the body must counteract these changes, which is done through the relaxation response. The relaxation response decreases heart rate, respiration rate and activity of the sympathetic nervous system through a reduction in the amount of cortisol and noradrenaline (Esch et al., 2003; Yehuda, 2011).

Music listening can promote the relaxation response by eliciting hormonal, immune, physiological, and cognitive changes (Iwanaga et al., 2005; Jacobs & Friedman, 2004; Khalfa et al., 2003; Knösche et al., 2005; Koelsch, 2005; Yamamoto et al., 2003; Yehuda, 2011). Studies found that listening to relaxing music decreased cortisol levels after stress (Khalfa et al., 2003) and decreased heart rate and respiration rate (Bernardi et al., 2006; de Witte et al., 2020). Research on brain responses during relaxation is limited and mostly focused on progressive muscle relaxation (PMR) and meditation. It is still unclear whether relaxation is associated with a decrease in brain activity, linked to lower heart rate and less engagement (Kobayashi & Koitabashi, 2016), or with an increase in brain activity, linked to increased focus and an active mental state (Lazar et al., 2000). During meditation, a meta-analysis found a global increase in attention-related areas and in somatosensory processing areas although the type of mediation (focused attention or mantras) and the duration of the experience impact brain responses (Tomasino et al., 2013). However, as this could be the specific effects due to meditation, more research on music-induced relaxation is warranted.

As music can have many effects on the brain and body (Brattico et al., 2011; Cohen, 2001; Eerola & Peltola, 2016; Frith, 2008; Herget, 2021; Huron, 2011; Jespersen et al., 2022), it is important to select the correct types of music. However, it is unclear which type of music should be used for relaxation. Physiological research suggests that music should have slow tempo and few dynamic changes to regulate breathing and heart rate (Bernardi et al., 2009, 2006; Esch et al., 2003; Merker et al., 2009). A meta-analysis found that most studies on music relaxation used soothing self-selected music which reduced stress and anxiety (Nilsson, 2008) however, other studies using different types of music are also effective (de Witte et al., 2020). It seems that the music that individuals choose themselves does not always fit the proposed calm music. Both calm music and more energetic music was found to be used by individuals in a sleep context (Dickson & Schubert, 2020; Scarratt, Heggli, Vuust, & Jespersen, 2023; Trahan et al., 2018). A similar dichotomy of music was found for self-selected music after stress (Adiasto et al., 2023). What individuals hope to achieve out of the music listening also impacts the features of the music (Adiasto et al., 2023; Scarratt, Stupacher, et al., submitted). For example, one study found that music to increase relaxation before sleep had less musical content whereas music to aid sleep through distraction had more musical content (Scarratt, Stupacher, et al., submitted). Given the many mechanisms that affect physiological responses to music, it is maybe necessary to broaden the scope of what type of music could be effective. For example, to act through episodic memory, it is necessary for the listener to have prior experiences with this music. In fact, if the music is presented repeatedly even energetic music could have a sedative effect (Iwanaga et al., 1996, 2005). Similarly, studies have found that hard rock music can increase participant relaxation ratings, although less than self-selected and soothing music (Burns et al., 2002; Labbé et al., 2007). Thus, how people respond to music is dependent on the music as well as their own past experiences with the music, their musical background, preferences, cultural and social factors, familiarity, and mood (Chew et al., 2016; Pereira et al., 2011; Tan et al., 2012; Yurdum et al., 2024).

While it is well researched that listening to the same music can elicit different emotions in individuals (Barrett et al., 2010; Juslin et al., 2016; Sung & Chang, 2005; Tan et al., 2012), how these individual differences are represented in the brain has often been overlooked. In beat and rhythm research, individual differences in beat perception has been found to correlate with activations in auditory and visual areas (Grahn & McAuley, 2009). In a neuroimaging study, both classical and R&B music showed similar brain area activations when the tracks were preferred by the listener (Wilkins et al., 2014). Similarly, personality traits, such as neuroticism and extraversion have been found to modulate neural responses to emotions expressed in music (Park et al., 2013). Investigating further the brain areas associated with different responses to music can clarify the involvement of musical characteristics and personal factors, like personality and musical familiarity.

Musical familiarity in particular can influence relaxation through various mechanisms (Chew et al., 2016; Finlay & Rogers, 2015; Freitas et al., 2018; Pereira et al., 2011). Firstly, musical familiarity may promote relaxation through decreasing the number of prediction errors and thus, alertness. Unexpected sudden changes in music increase arousal (Huron, 2006; Yehuda, 2011). If a track is familiar to the listener, they have more precise expectations and less prediction errors, leading the changes to be less salient, resulting in less alertness and more relaxation (Koelsch et al., 2019; Vuust et al., 2022). Secondly, listening to familiar music might trigger memories (Gerdner, 2000; Hicks-Moore, 2005; Juslin & Västfjäll, 2008; Sung & Chang, 2005). Finally, the memories or the music itself might induce pleasure, and an increased activation of the limbic system, increasing relaxation through pleasure (Pereira et al., 2011; Peretz et al., 2009). A meta-analysis investigating the neural correlates of musical familiarity reported the left superior frontal gyrus, the ventral lateral nucleus of the left thalamus, and the left medial surface of the superior frontal gyrus as being the most important for familiar music and the left insula and right anterior cingulate cortex for unfamiliar music. Familiar music was associated with a motor pattern of activation, reflecting an audio-motor synchronization to the rhythm and/or a covert sing-along response (Freitas et al., 2018).

In this paper, we investigate the perceived effect of music on relaxation, in particular the influence of musical familiarity and musical features and its neural correlates. The musical features in this study are referred to as Type and include Excitative music, energetic music, and Sedative music, calm music. We hypothesise that listening to Sedative music will provide stronger relaxation responses than Excitative music. Similarly, we expect Familiar music to provide stronger relaxation responses than Unfamiliar music. We also hypothesise that Familiar music will have more activation in memory-, motor-, and emotion-related areas. Contrastingly, listening to Unfamiliar music will activate attention-related areas and Excitative music will activate auditory-motor and attention-related areas.

## Methods

### Stimuli

This project used 28-second excerpts of 48 songs from a newly created dataset, the Relaxation Music Dataset, collected following the methods described in Scarratt, Heggli, Vuust, & Sadakata (2023) and in Heggli, (2020). To form this dataset, we automatically collected all playlists on Spotify with ‘Relax’ in the title or description. In total, 1,245 playlists were identified. The audio features and information of the tracks from these playlists were extracted. There were 321,866 tracks in total of which 187,842 unique tracks. The unique tracks were then split on the Spotify features *Energy* and *Popularity* to select 96 tracks balanced on these features. The unfamiliar tracks were matched to the familiar tracks based on genre, style, and release date. After a preliminary pilot study to verify familiarity of these tracks, a final 48 tracks were selected (Supplementary Table S32), with 12 per condition in a 2×2 design Familiarity (Familiar or Unfamiliar) and Type (Sedative: slow, calm music or Energetic: upbeat music). After selection, the tracks were cut to 28-seconds and normalised making sure that the most recognisable section of the melody was selected for the familiar tracks. This duration was chosen as it was the maximum length to be able to repeat all four conditions within the 128 Hz of the magnetic resonance imaging (MRI) scanner. Any longer duration would lead to drift errors in the results.

### Participants

Participants were recruited through the university’s network and the local Sona recruitment system. Participants were included if they were healthy, comfortable speaking and reading English and had no MRI-contraindications. To have participants with a comparable music exposure, participants younger than 19 and older than 40 were excluded. 58 participants matched the inclusion criteria and completed the scan. The remaining 58 participants had a mean age of 25.55(4.61) and an age range of 19-40. There were 40 women, 17 men and one person who identified as non-binary. Most participants were living in Denmark at the time of the study (51) although only a few were Danish (11). Through a self-reported question, 52 participants indicated that they were right-handed, 2 were left-handed and 4 were ambidextrous. The highest completed education for 17 participants was high school, 22 participants had received a bachelor’s degree, 17 had received a master’s degrees, one had completed their PhD, and 1 participant had stayed in education longer. One participant (F, 26 years) was excluded from the behavioural and clustering analyses due to too many missing values but was included in the overall neuroimaging analyses. Characteristics of participants can be found in Supplementary Table S1.

### Procedure

Before coming into the lab, participants were asked to select 12 out of 18 of the Excitative Familiar and 12 out of 18 of the Sedative Familiar tracks. This step was to verify their familiarity with the selected tracks. Once in the lab, participants were explained the MRI procedure, signed the informed consent and MRI safety forms. The MRI session consisted of a 10-min resting state scan, followed by two blocks of 14 minutes of the main task. In each block, they heard in a pseudorandom order 6 tracks from each of the four music conditions (Familiar-Sedative, Familiar-Excitative, Unfamiliar-Sedative, Unfamiliar-Excitative). The order of presentation was the same within a block but randomly assigned and randomly different between both blocks. After listening to each 28-second track, participants were asked to rate how relaxed they felt on a scale of 0-100. They were instructed to try to focus on how their body was reacting rather than how they thought they should be feeling, to get an accurate measure of their relaxation response and not their cognitive judgment of the music (Scarratt, Labouriau, et al., submitted). Finally, there was a 10-min anatomical scan. After the MRI session, participants were asked to complete a set of questionnaires. This included demographics, musical background, and the extraversion subscale of the Eysenck personality test as extraversion can impact music preference (see Supplementary Table S2). Participants also listened to the tracks once more for 5-seconds and rated how familiar they were with the track and how much they liked it. In total, the MRI session took 1h. Preparation and the questionnaires took 1h.

Participants were compensated for the two hours of their participation. The project received ethical approval from the Institutional Review Board at DNC (DNC-IRB-2023-005) and was conducted according to the *Helsinki Declaration II*.

### MRI acquisition

Using a Siemens Prisma 3.0T MRI scanner, we collected 820 volumes during the tasks and 600 volumes during resting state using echo-planar imaging (EPI) with a multi band sequence (TR=1000ms, TE=29.6ms, voxel size=2.53*2.53*2.50mm, flip angle=60 degrees, slice thickness=2.5mm, acceleration factor = 3, 2.5mm spacing between slices, 54 slices, matrix=76*76 and field of view (FOV) = 1535*1535 mm). For spatial normalization and localization, a high-resolution T1-weighted anatomical image was then acquired using a magnetization prepared gradient echo sequence (MP2RAGE, TR=5000ms, TE=2.87ms, TI1=700ms, TI2=2500ms, voxel size=0.90*0.90*0.90mm, flip angle 1=4 degrees, flip angle 2=5 degrees, slice thickness=0.9mm, 192 slices, matrix=256*240 and FOV = 215*230 mm).

The music was presented to participants with OptoActive™ II MR compatible headphones (Optoacoustics Ltd, Mazor, Israël). Participants wore in ear earplugs to protect their ears from the noise of the scanner, and the music’s volume was adjusted to each individual’s comfort level.

### Behavioural Analyses

#### Reclassification of familiarity

Although we piloted the familiarity of the tracks and asked participants to select the 12 tracks that they were most familiar with, we also asked participants to rate their familiarity with each track outside of the scanner. In a few cases, their rated familiarity with a track did not correspond to how it was classified. Therefore, we reclassified these tracks per participant based on their actual rated familiarity. We separated the data based on whether it was initially categorised as Familiar or Unfamiliar and calculated the mean and standard deviation for each of these. A trial was recategorized if the measured familiarity was higher or lower than the mean for that category plus or minus two standard deviations of that category. This means that any familiar-categorised trial that had a measured familiarity less than 46 (Mean of 86.56 – 2* sd of 20.33) was reclassified as Unfamiliar. Any unfamiliar-categorised trial that had a measured familiarity of more than 68 (mean of 19.57 + 2* sd of 24.23) was reclassified as Familiar. 85 tracks were reclassified as Unfamiliar, and 79 tracks were reclassified as Familiar. We reclassified at least one trial for 42 participants, giving an average of 2.78 trials reclassified per participant.

The relaxation responses were modelled in a Bayesian way. Relaxation and Liking ratings were rescaled from 0-100 to 0-1 and Liking was z-scored. The relaxation ratings followed a beta zero-one inflated distribution. We ran multiple stepwise models using the variables Type, Familiarity, Liking and the random effects of participant for Type and Familiarity. We also fitted simple beta regression models by changing any relaxation 0 value to 0.0001 and any 1 value to 0.9999. We then fitted multiple simple models stepwise adding Type, Familiarity, Liking and random effects. All estimates for the tested models are in Supplementary Tables S3-S10.

When comparing the models with Watanabe-Akaike information criteria (WAIC), p_waic and predictive accuracy (ELPD_waic), Supplementary Table S11, the model that had a good balance of predictive accuracy, best model fit and acceptable complexity was Relaxation ∼ Type + Familiarity + Liking + (1 + Type + Familiarity | ID, so we proceeded further with that model. However, it is important to note that the estimates for Sedative, Unfamiliar and Liking remained relatively similar across the various models, meaning that the results are robust, and the choice of model does not impact the results.

All Bayesian analyses were done using the RStudio software and the brms package.

Additionally, for more robustness, we performed the same analysis using linear regression with a beta distribution using the R package GlmmTMB and found similar results (Supplementary Tables S12-S22 and Supplementary Analysis S1).

#### Clustering

The spread and weight of Type and Familiarity was very different depending on the individuals (Supplementary Figure S1) To explore this individual variation, we performed a clustering analysis on the inverse-logit random effects of each participant added to the fixed effects. To have robust clusters, we did three different clustering analyses and summarised the findings to get the most consistent results. We first split the data into 4 clusters based on whether the estimated effects per participant were positive or negative for Type and Familiarity (Supplementary Figure S2). We also did a k-means clustering analysis on the same data (Supplementary Figure S3). As we would be using the results of the clustering analysis for further MRI analyses, the clusters needed to not be too small.

Therefore, we also performed a balanced k-clustering analysis (Supplementary Figure S4). The results from each were summarised into the final clustering we used (Supplementary Table S23 and Supplementary Table S5).

All behavioural data and analysis scripts can be found at GitHub/RebeccaJaneScarratt/MusicRelax

### fMRI Analysis

#### Preprocessing

The functional images were pre-processed and analysed using SPM12 (http://www.fil.ion.ucl.ac.uk/spm) running on Matlab R2016b (MathsWorks, MA, USA). The origins of each session were reset manually to the anterior commissure. The ‘Realign and Unwrap’ function in SPM (Statistical Parametric Mapping) was employed to correct for head motion and associated intensity variations in fMRI images. Initially, the first image of every session for each participant was aligned to the first image of the initial session to ensure consistent session alignment. Within each session, all subsequent scans were then realigned to the first image of that session, ensuring intra-session consistency. After realignment, physiological noise was managed using the FIACH-package (Tierney et al., 2016), which identifies and corrects non-physiological large amplitude temporal signal changes and regions of high temporal instability. Noisy voxels in the realigned EPI images were interpolated using a natural cubic spline, utilizing the two adjacent time points. Subsequently, the structural scans were coregistered to the mean functional image derived from the realignment step. The unified segmentation model (Ashburner & Friston, 2005) was then applied to the structural image to segment grey and white matter as well as cerebro-spinal fluid, which included bias field correction. The derived deformation fields were used to normalize the FIACH-filtered functional images to the standard Montreal Neurological Institute (MNI) stereotactic space. Finally, the images were smoothed with an 8 mm full-width at half-maximum (FWHM) Gaussian kernel to enhance the signal-to-noise ratio for subsequent statistical analysis.

Movement artifacts were further mitigated by applying the Artifact Detection Tool (ART– RRID:SCR_005994) to identify movement outliers based on excessive framewise displacement (FD). Physiological data were processed using a custom Python script designed to parse DICOM files and extract heart rate and respiration signals. The script applies peak detection and filtering to generate noise regressors based on RETROICOR modelling (RETROspective Image-based Correction of physiological artifacts; Glover et al., 2000), aligning physiological fluctuations with fMRI data.

#### Statistical Analysis

Individual statistical maps (first-level analysis) were generated for each subject, with separate regressors for each condition. To account for residual motion and noise-related artifacts, movement regressors from realignment, noise components identified using FIACH, RETROICOR physiological noise regressors, and outlier time point detected with ART were included as covariates in the first-level analysis.

These results were combined in full factorial general linear model second-level analysis, with two factors each with two levels: Familiarity (familiar-unfamiliar) x Type (sedative-excitative). A grey matter mask was defined using the AAL3 atlas, excluding visual areas (Rolls et al., 2020; Tzourio-Mazoyer et al., 2002; Supplementary Table S24). Main effects of Familiarity and Type were obtained using familywise error (FWE) correction for multiple comparisons with a p value of p = .05 and cluster size k>10. *T-tests* were then performed on the defined contrasts of interest (Familiar vs Unfamiliar, Unfamiliar vs Familiar, Sedative vs Excitative and Excitative vs Sedative).

Subsequently, another second-level analysis was performed based on behavioural clustering results. One factorial design model (Familiarity x Type) was performed per cluster, the grey matter mask described above and main effects of Familiarity and Type and *t-tests* for each contrast. All results were familywise error corrected with a p-value of p=.05 and a cluster size k>10. All anatomical labelling of clusters was based on the probabilistic neuromorphometrics atlas (https://www.neuromorphometrics.com) within SPM.

## Results

### Sedative music followed by Familiar music is most relaxing

57 participants rated how relaxed they felt listening to 28-second excerpts of Sedative or Excitative and Familiar or Unfamiliar tracks. They rated tracks in all conditions with an average relaxation of 59.85(24.54). For the condition Excitative-Familiar had relaxation ratings of 60.79(26.77), the condition Sedative-Familiar had relaxation ratings with a mean of 70.32(22.22). The average relaxation ratings for condition Excitative-Unfamiliar were 49.65(21.95) and for the condition Sedative-Unfamiliar, relaxation ratings were on average 58.54(22.38; Figure 1A).

**Figure 1:**
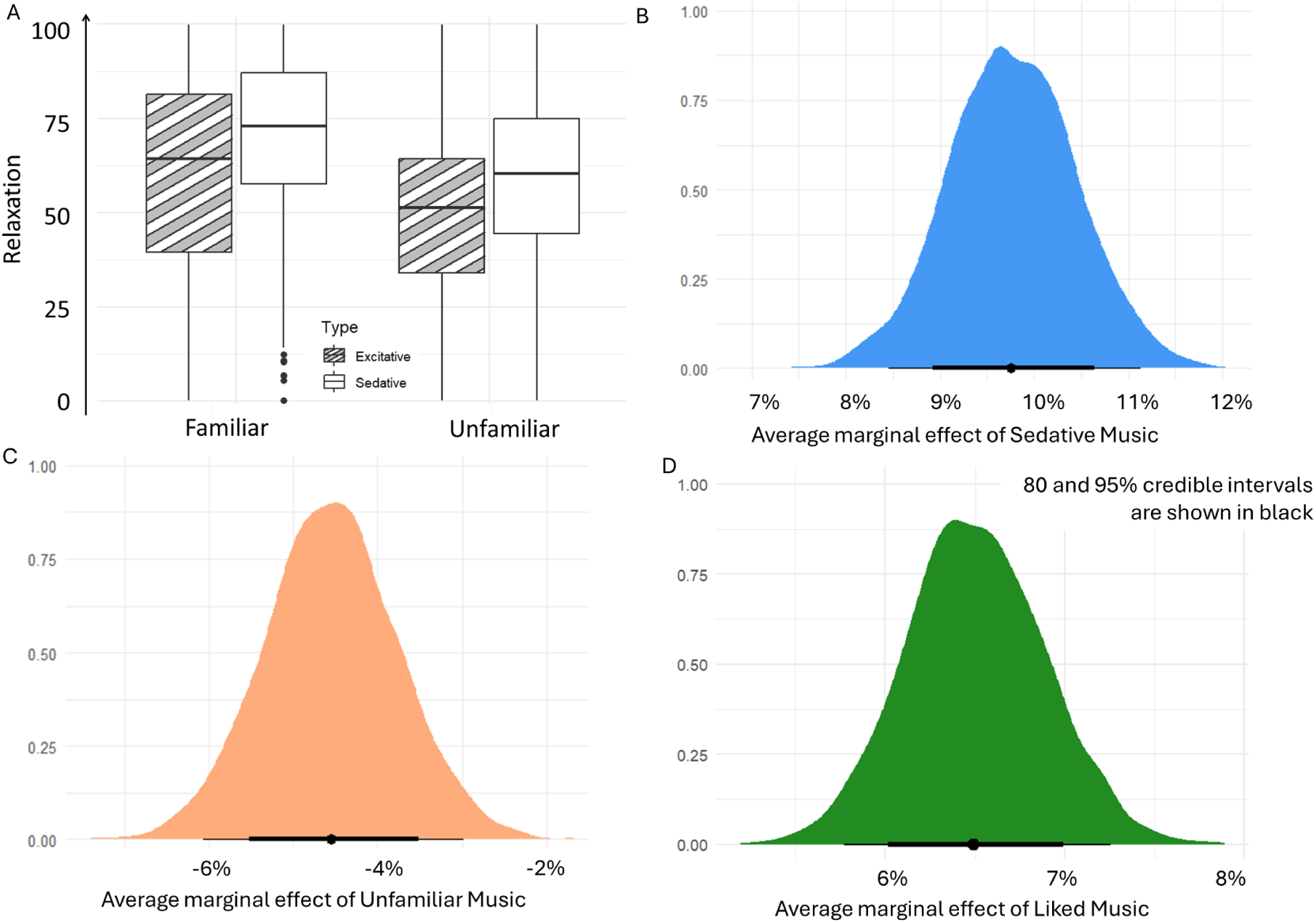
Relaxation responses per condition. *A)* shows the plotted raw data of relaxation ratings per Familiarity and Type. The results showed that Sedative-Familiar music had the highest relaxation scores whereas Excitative-Unfamiliar had the lowest. *B)* The Bayesian analysis showed that the Type of music had the largest impact on relaxation ratings which increased with 9.8% when moving from Excitative to Sedative music, C) Similarly, unfamiliar music decreased relaxation levels by -4.6% and D) liking the music increased relaxation ratings by 6.5%.

The Bayesian analysis of the relaxation ratings revealed that on average with Excitative and Familiar music, Relaxation is at 65%. Having Sedative music increased Relaxation by about 9.8% with a 95% credible interval of 8.4%-11.1%, having unfamiliar music decreased Relaxation by about 4.6% with a credible interval of -6.1% to -3.0% and liking the music increased Relaxation by 6.5% with a credible interval of 5.75-7.26 (Figure 1 B, C, D).

### Patterns of brain activity for Familiarity and Type of music

We are interested in how musical familiarity and musical features influence brain activity. The significant activations for the main effect of Familiarity, Type and their Interaction are in Supplementary Table S25. The significant activations from the contrasts Familiar vs Unfamiliar, Unfamiliar vs Familiar and Excitative vs Sedative are in Supplementary Table S26. There were no significant peaks after correcting for family-wise error for Sedative vs Excitative. Details of the significant activations are described below.

#### Excitative vs Sedative Music

Looking at the main effect of Type on brain activity by comparing Excitative with Sedative music, we see increased activation in auditory areas: bilateral Heschl’s gyrus and right planum polare, and motor areas: left Cerebellum, left superior frontal gyrus (SFG), right supplementary motor area (SMA), right putamen, and left thalamus (see Figure 2 and Supplementary Table S25). Similarly, post-hoc contrast ‘Excitative vs Sedative’ reveals activity in primary and secondary auditory areas: bilateral Heschl’s gyrus, transverse temporal gyrus (TTG), and BA22; and motor areas: bilateral cerebellum, bilateral SMA, bilateral putamen, right caudate, and left thalamus (see Figure 2 and Supplementary Table S26).

**Figure 2:**
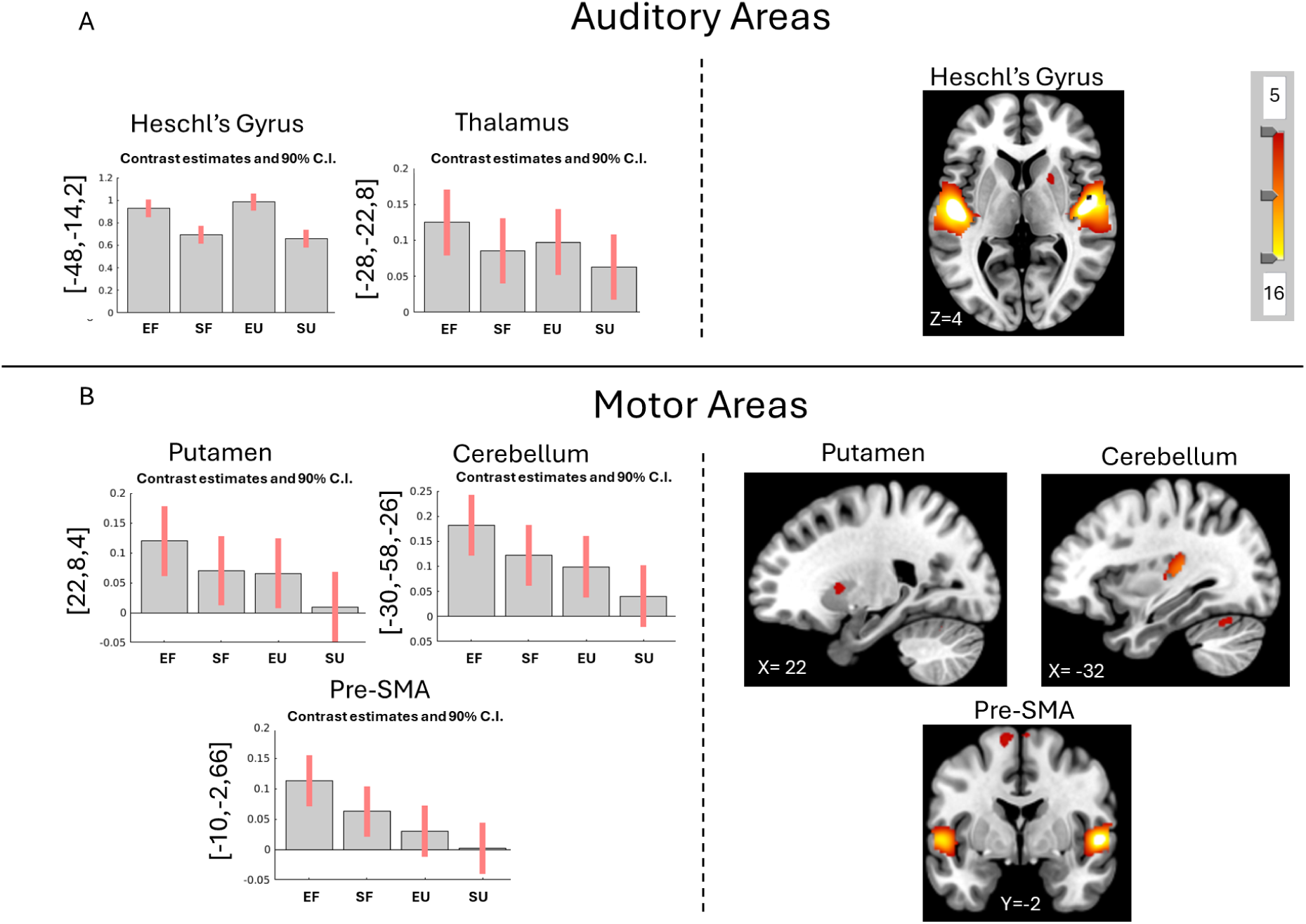
Main effect of Type and Contrast Excitative vs Sedative for all participants. Main effect of type (bar plots) reveals more activity (A) auditory (bilateral Heschl’s gyrus, left thalamus) and (B) motor areas (left cerebellum, left SMA, and right putamen). Similarly, the pos-hoc contrast Excitative vs Sedative music (images) reveals increased activity in (A) auditory (bilateral Heschl’s gyrus) and (B) motor areas (bilateral cerebellum and bilateral putamen). More details and coordinates can be found in Supplementary Tables S25 and S26. FWE p<.05, k>10. Coordinates are presented in MNI. Abbreviations: SMA= supplementary motor area, EF= Excitative-Familiar, EU= Excitative Unfamiliar, SF= Sedative Familiar, SU = Sedative Unfamiliar

#### Familiar vs Unfamiliar Music

The main effect of Familiarity on brain activity revealed increased activity when listening to familiar music in auditory areas: left superior temporal gyrus (STG), right middle temporale gyrus (MTG), left inferior frontal gyrus (IFG), bilateral temporal pole, and right planum temporale; in motor areas: bilateral precentral gyrus, bilateral cerebellum, right pre-SMA, and left putamen; in memory areas: posterior dorsal cingulate gyrus; and in emotion areas: left amygdala (Figure 3 and Supplementary Table S25).

**Figure 3:**
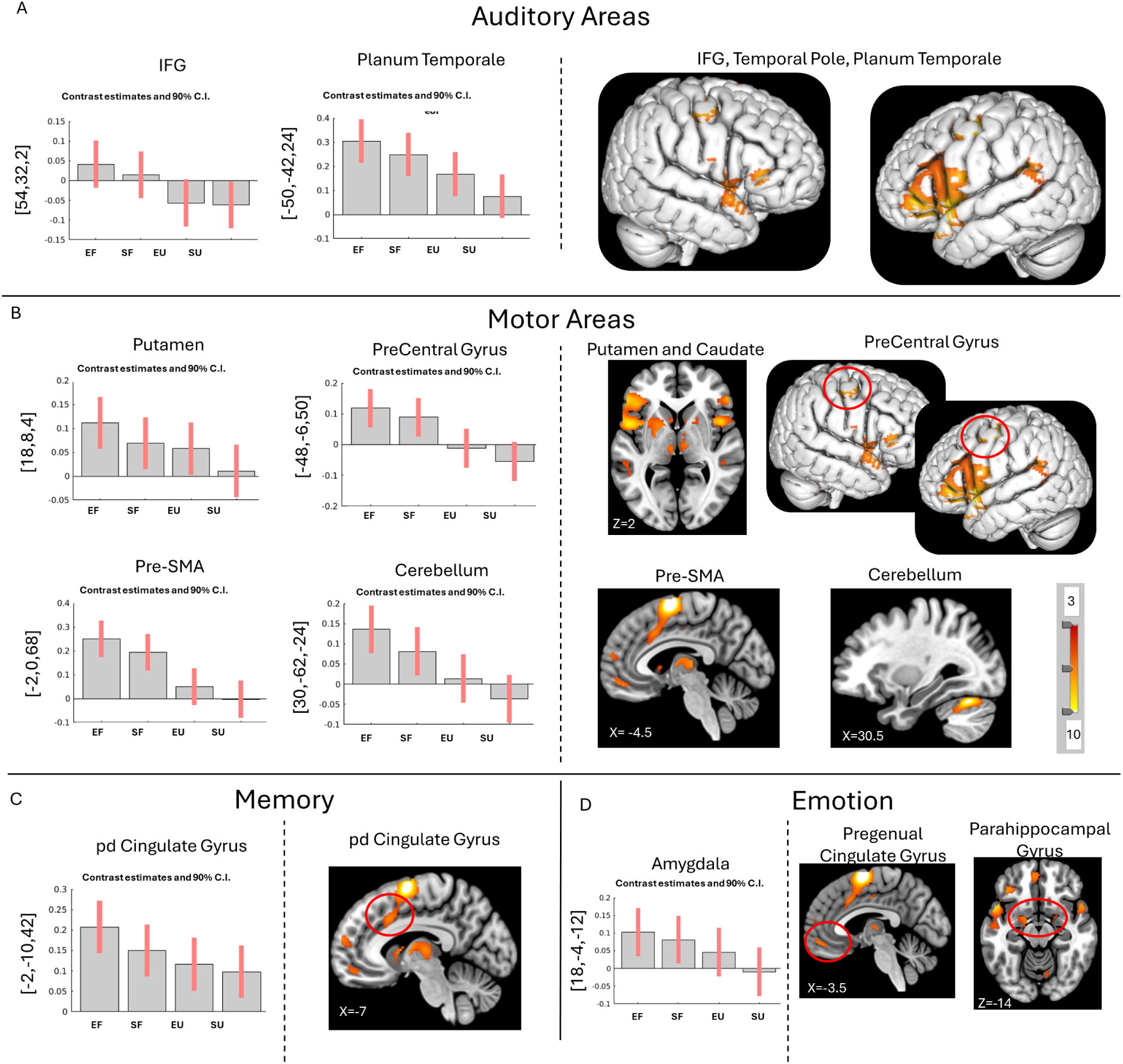
Main effect of Familiarity and Contrasts Familiar vs Unfamiliar over all participants. Main effect of familiarity (bar plots) reveals more activity in areas linked with (A) auditory (left IFG, bilateral temporal pole, right planum temporale), (B) motor (left putamen, bilateral precentral gyrus, bilateral cerebellum, right pre-SMA), (C) memory (right cingulate gyrus) and (D) emotion (amygdala) processing. The contrast Familiar vs Unfamiliar (images) music reveals increased activity in (A) auditory (bilateral SFG including temporal pole and planum temporale, bilateral IFG), (B) motor (bilateral cerebellum, left putamen, right pre-SMA, bilateral precentral gyrus), (C) memory (right posterior dorsal cingulate gyrus) and (D) emotion (right pregenual cingulate gyrus and right parahippocampal gyrus). More details and coordinates can be found in Supplementary Tables S25 and S26. FWE p<.05, k>10. Coordinates are presented in MNI. Abbreviations: IFG=inferior frontal gyrus; SFG=superior frontal gyrus; SMA= supplementary motor area, pd= posterior dorsal, EF= Excitative-Familiar, EU= Excitative Unfamiliar, SF= Sedative Familiar, SU = Sedative Unfamiliar

Post-hoc contrast analysis of Familiar vs Unfamiliar music revealed increased activation in auditory areas: bilateral Heschl’s gyrus, left temporale pole, bilateral BA22 and right BA13; motor areas, right precentral gyrus, bilateral cerebellum, and left putamen; a memory-related area: right posterior dorsal cingulate gyrus; emotion-related areas: right pregenual cingulate gyrus and right parahippocampal gyrus (Figure 3 and Supplementary Table S26).

#### Unfamiliar vs Familiar Music

When listening to unfamiliar music, the main effect reveals increased activity in attention-related areas: left MFG and left angular gyrus (Figure 4 and Supplementary Table S25).

**Figure 4:**
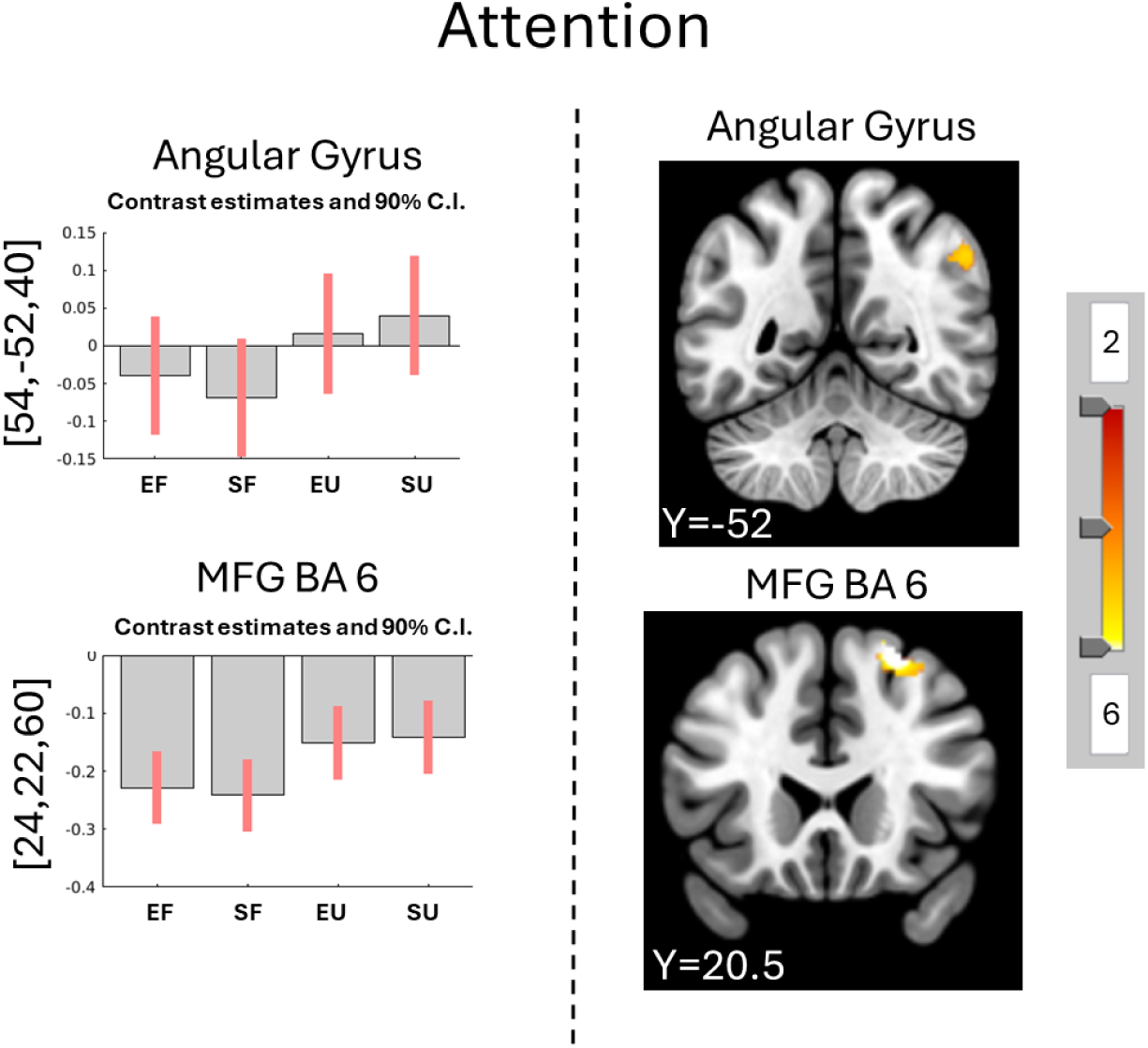
Main effect of Familiarity and the Contrast Unfamiliar vs Familiar music over all participants. The main effect of Familiarity reveals more activity in the angular gyrus and MFG for unfamiliar compared to familiar music (bar plots). The contrast Unfamiliar vs Familiar music finds the same areas more involved in processing Unfamiliar compared to Familiar music (images). More details and coordinates can be found in Supplementary Tables S25 and S26. FWE p<.05, k>10. Coordinates are presented in MNI. Abbreviations: MFG= middle frontal gyrus, EF= Excitative-Familiar, EU= Excitative Unfamiliar, SF= Sedative Familiar, SU = Sedative Unfamiliar

Post-hoc contrast analysis of Unfamiliar vs Familiar revealed increased activity in attention-related areas: left MFG BA8, left angular gyrus, and left BA10 (Figure 4 and Supplementary Table S26).

### Four distinct clusters of relaxation responses to music

To further explore the individual differences in relaxation rating in response to music, we clustered the relaxation ratings. Using the compound clustering analysis described above, we found four distinct clusters that showed different relaxation responses to Type and Familiarity (Figure 5 and Supplementary Figure S5 and Supplementary Table S23).

**Figure 5:**
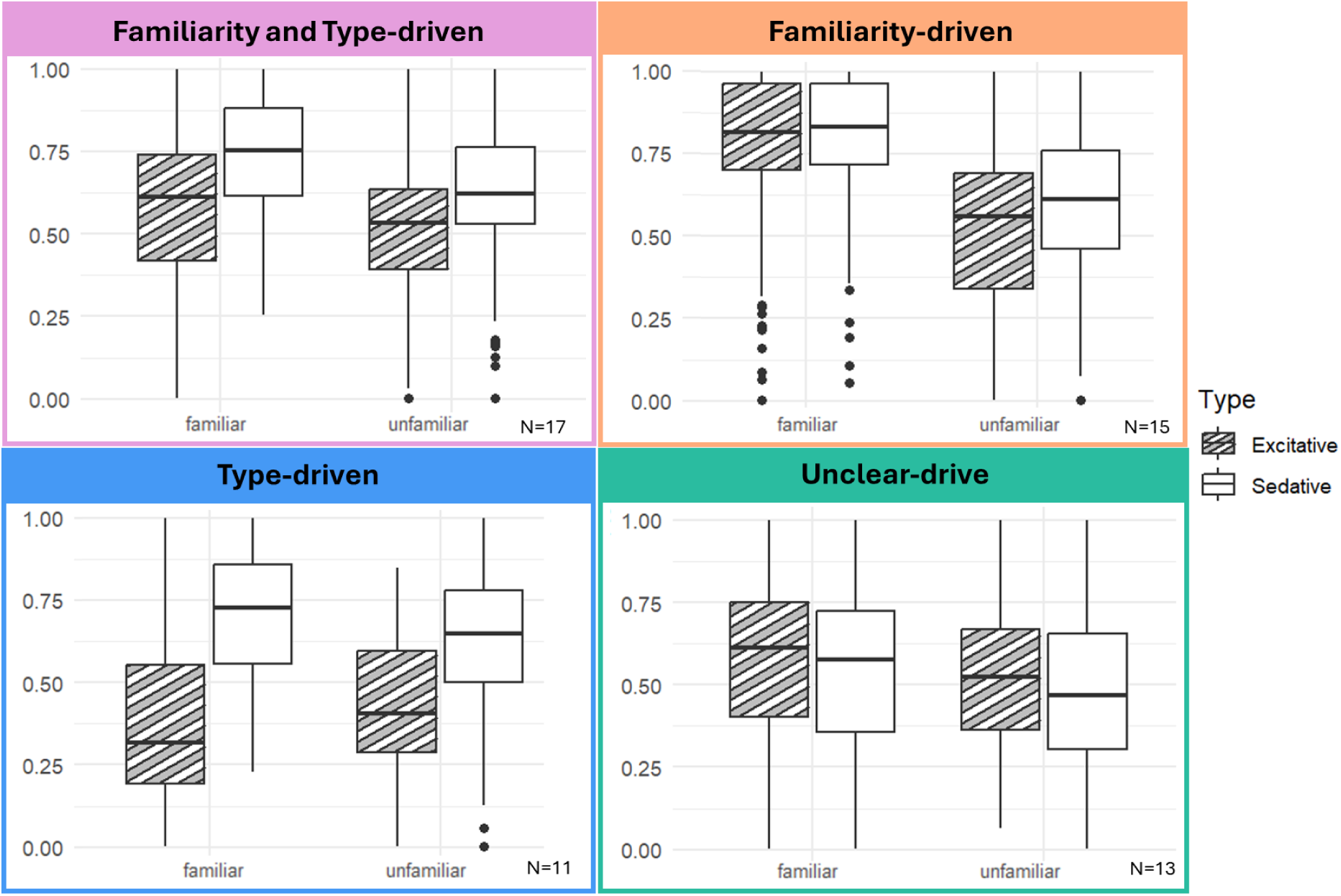
Relaxation by Type and Familiarity for each of the four identified clusters. Through a combination of clustering methods on the random effects of Familiarity and Type per participants, we identified four subgroups who rated relaxation differently to these conditions. One group responded as the average (Familiarity and Type-driven), one group mostly responded to Familiarity (Familiarity-driven), one group mainly responded to Type (Type-driven), and the last group had unclear patterns (Unclear-drive).

We named the clusters based on the factors that seemed to drive each cluster the most. The first cluster responded the most like the overall findings which are exactly as predicted so was named “Familiarity and Type-driven”. The second cluster clearly had the highest relaxation ratings for familiar music therefore was named “Familiarity-driven”. The third cluster had the biggest difference between Excitative and Sedative music so was named “Type-driven”, and the final cluster had a small difference in that familiarity was higher than unfamiliar, but they responded in an opposite way to Type with Excitative music being higher than Sedative music, so it was named “Unclear-drive”. Demographic information of participants in each cluster is reported in Supplementary Table S31.

### Four distinct brain activity patterns

We then performed a GLM for the fMRI data within each of these clusters. The results of the main effects and contrasts for each cluster are in Supplementary Tables S27-S30.

#### Excitative vs Sedative Music

When comparing within each cluster the effect of Type on brain activity, we found that all clusters had more activity in the auditory cortex for Excitative vs Sedative music, similarly to the main results. More specifically, Type-driven had activity focused on Heschl’s gyrus whereas the other clusters had more peripheral auditory cortex activations (Figure 6 and Supplementary Tables S27-S30).

**Figure 6:**
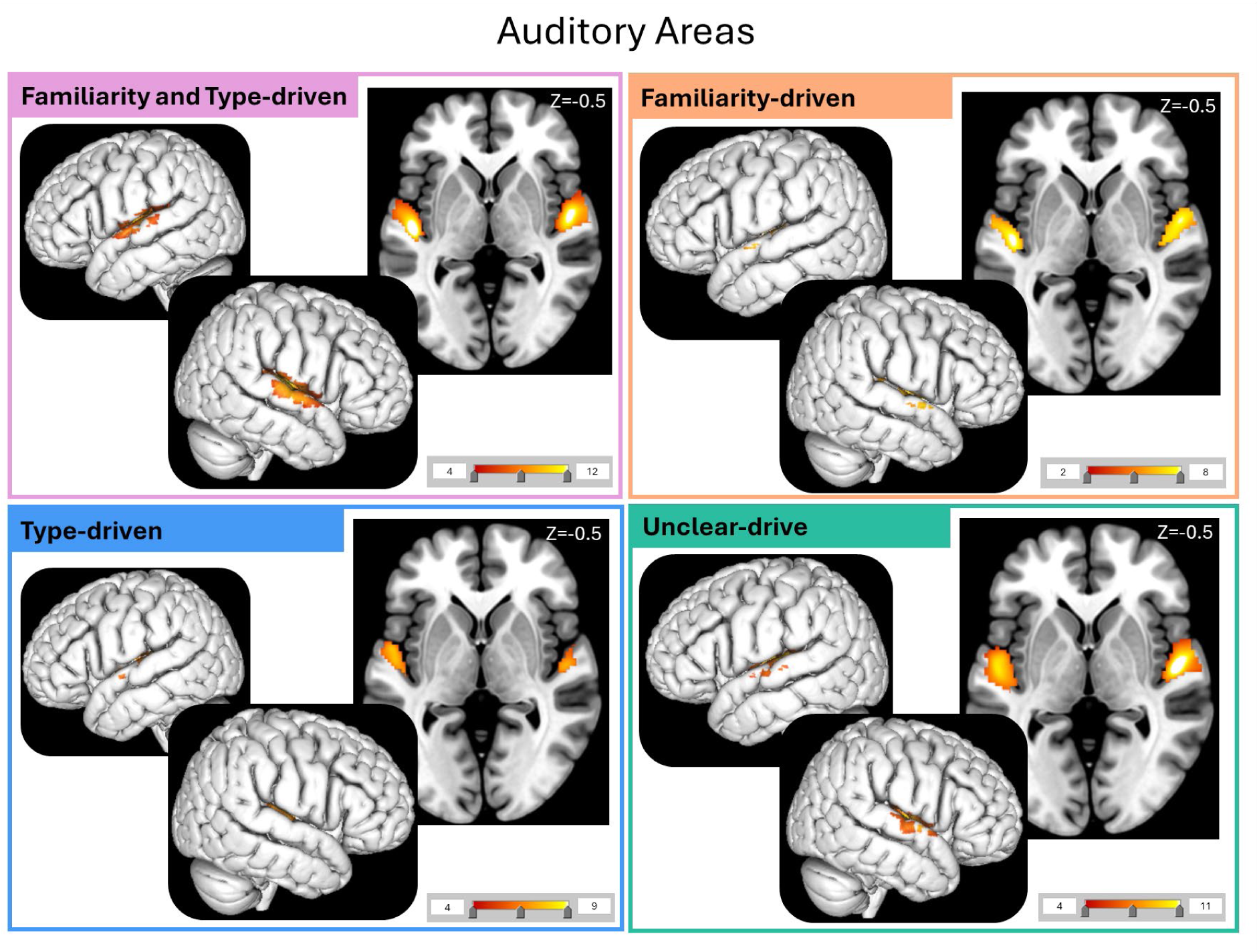
Auditory Areas activated by the Contrast Excitative vs Sedative music per cluster. The four analyses reveal differences in activation in auditory areas between the four clusters. Namely, the auditory activations in the Familiarity and Type-driven and Unclear-drive clusters had more peripheral activation compared to more central Heschl’s gyrus activation in the Type-driven cluster. The cluster Familiarity-driven cluster showed some peripheral activations but not as much as the previously mentioned clusters.

#### Familiar vs Unfamiliar Music

When comparing within each cluster the effect of Familiarity on brain activity, we found that only the cluster Unclear-drive had more activity for familiar music compared to unfamiliar music in the auditory cortex (Figure 7A and Supplementary Tables S27-S30). We found that all clusters except for the Type-driven cluster had more activity in the motor cortex for Familiar vs Unfamiliar music. More specifically, the Unclear-drive, Familiarity-driven, and Familiarity and Type-driven clusters had more activity in the pre-SMA, precentral gyrus and cerebellum for Familiar music. The Familiarity-driven cluster also had more activity in the thalamus (Figure 7B and Supplementary Tables S27-S30). The clusters Familiarity-driven and Type-driven also had activation in the inferior frontal gyrus (IFG) and Operculum 9/10, respectively (Figure 7C and Supplementary Tables S27-S30).

**Figure 7:**
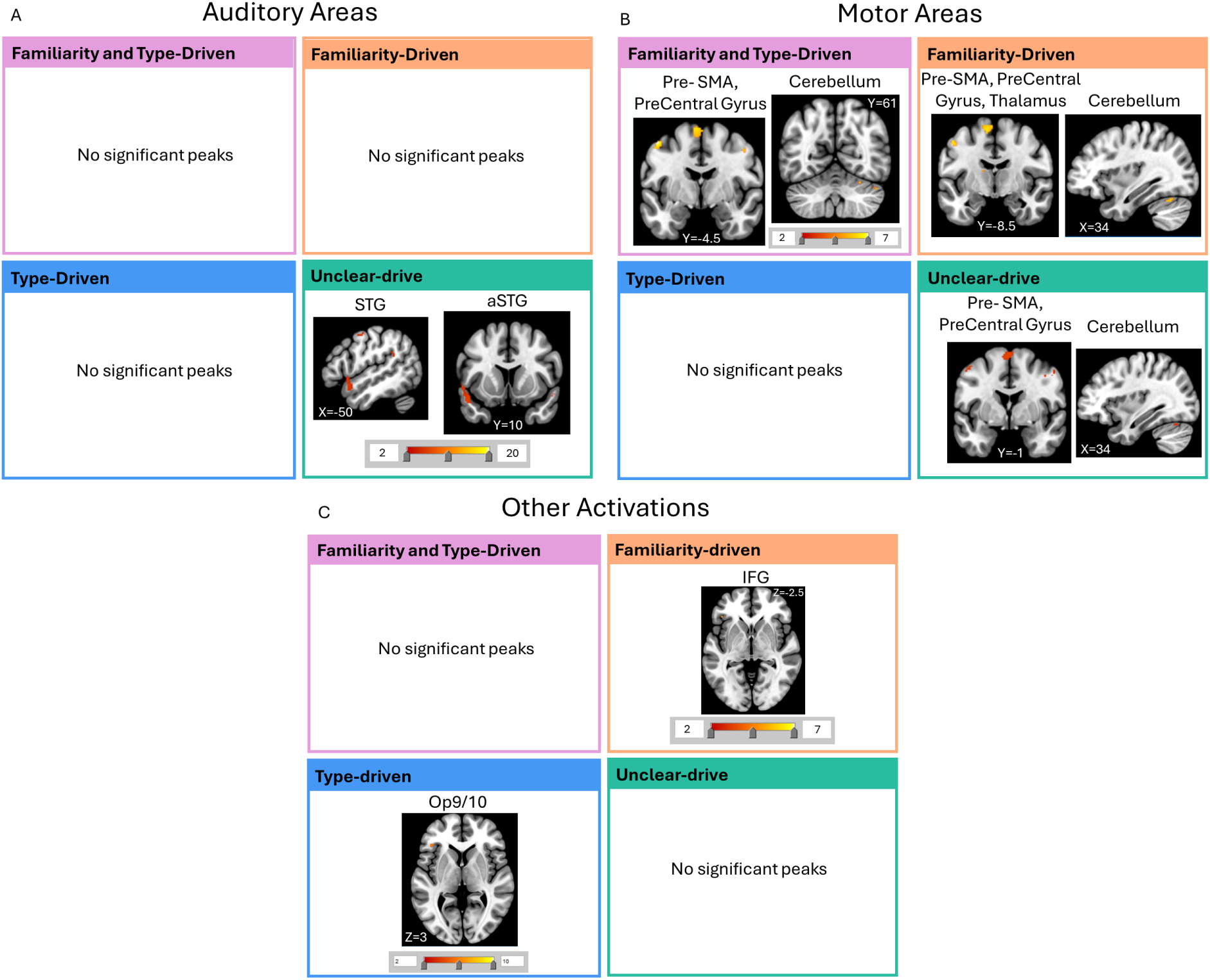
Auditory, Motor and Other Areas activated by the Contrast Familiar vs Unfamiliar music per cluster. A) The four analyses revealed more auditory activation for the cluster Unclear-drive in the aSTG and STG. B) There was more activity in the pre-SMA, precentral gyrus, and cerebellum for all clusters except Type-driven. There was also more activity in the Thalamus and IFG for Familiarity-driven cluster. C) Finally, there was activation of the Operculum in Type-driven. Abbreviations: IFG= inferior frontal gyrus, STG= superior temporal gyrus, SMA= supplementary motor area, Op= Operculum

#### Unfamiliar vs Familiar Music

Unfamiliar music led to more activity in the MFG compared to Familiar music for Familiarity-driven cluster (Figure 8 and Supplementary Tables S27-S30).

**Figure 8:**
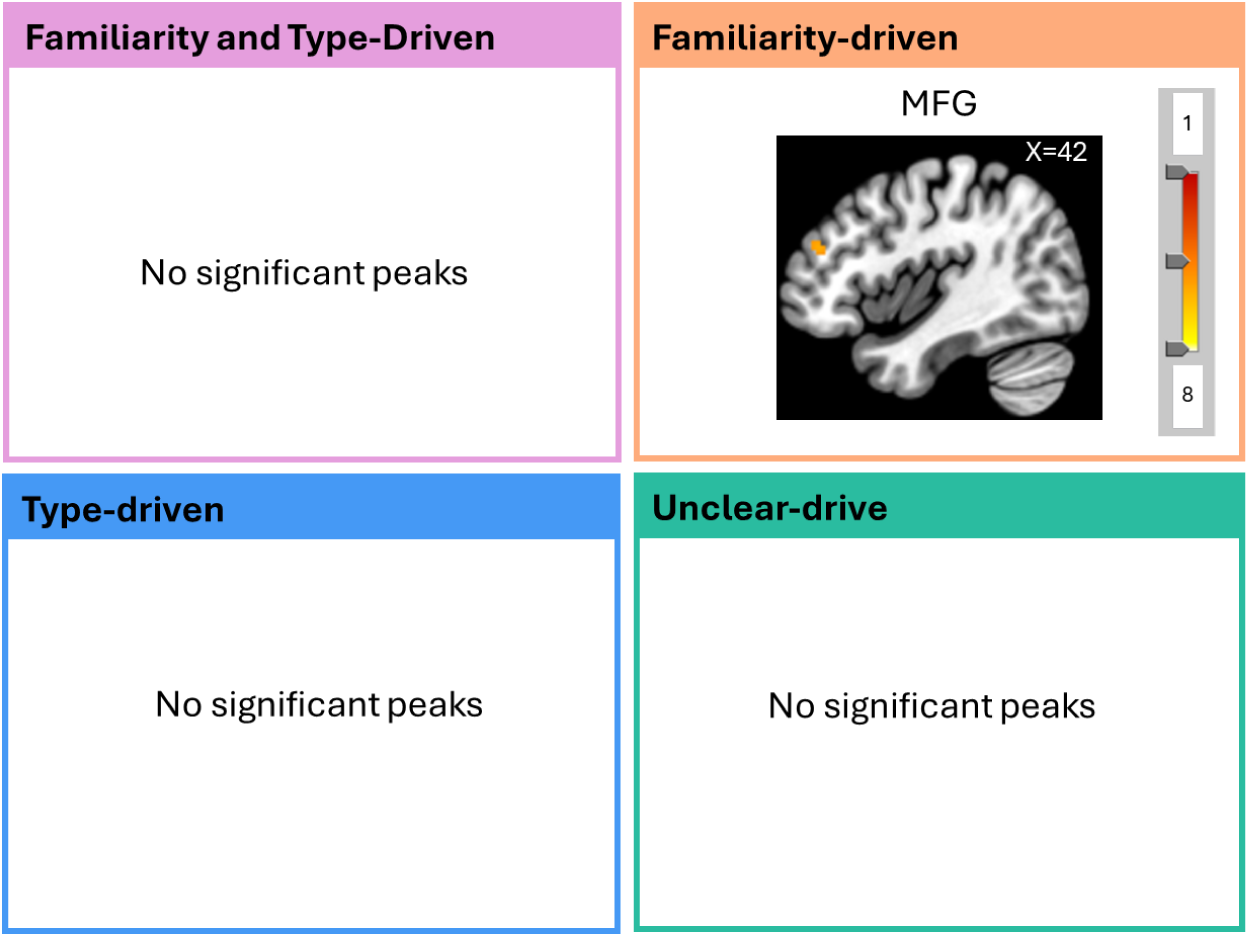
Regions activated by the Contrast Unfamiliar vs Familiar music per cluster. The contrast unfamiliar vs familiar music only revealed significant activations for the Familiarity-driven cluster where the MFG was more involved. Abbreviations: MFG=middle frontal gyrus

## Discussion

This paper shows that different neural activations underlie the processing of different types of music used for relaxation. On the behavioural level, having Sedative and Familiar music increases relaxation compared to Unfamiliar and Excitative music. These behavioural findings are consistent with neuroimaging results which show increased auditory, motor and cognitive processing for Familiar music, more attention processing for Unfamiliar music, and increased auditory processing for Excitative music. Additionally, we see evidence that the same music does not affect individuals in the same way. We find four distinct clusters for which Familiarity and Type influence relaxation differently. The fMRI analysis of these clusters reveals that key differences between the groups are the way they process auditory information, whether they engage motor areas, and how much attention areas are recruited.

### Music for Relaxation: Effect of Music Type and Familiarity

The behavioural results of this study show that how relaxed an individual feels when listening to music is influenced by the content of the music, with more relaxation for Familiar, Sedative, and liked music, as was hypothesised. These effects are linked to neurological differences. Increased attention is needed to process Unfamiliar music, emotions are involved in Familiar tracks, and more auditory processing is necessary for Excitative tracks.

Listening to Excitative music led to more activations of auditory networks, specifically in Heschl’s gyrus and the thalamus. This is in line with the fact that Excitative tracks had more musical content and energy, therefore had more auditory information to process. Excitative music also led to more activation of motor areas, such as the putamen, cerebellum and pre-supplementary motor area (SMA) than Sedative music. Previous studies have shown motor activation when just listening to music, especially when the music is groovy and has a strong rhythmic component (Matthews et al., 2020; Zalta et al., 2024; Zatorre et al., 2007). This motor activation could also be linked to participants engaging more with the music and/or covertly singing along to lyrics. A meta-analysis found these exact areas to be important for singing, both covertly and overtly (Scarratt et al., in preparation). Alternatively, putamen activity could be related to relaxation as meditating is linked to increased putamen activation (Lazar et al., 2000).

Behaviourally, Familiar and liked music was positively associated with relaxation, although slightly less than Sedative music. Listening to Familiar music activated the auditory network, although mostly peripheral auditory areas such as the inferior frontal gyrus (IFG), planum temporale and temporal pole, indicating that the auditory stimuli of Familiar music was processed at a higher level, possibly relating to lyric comprehension or prediction (Binder et al., 1996; Brugge et al., 2003; Foundas et al., 1994; Krumbholz et al., 2005). Motor areas such as the putamen, caudate, cerebellum, pre-SMA and precentral gyrus were also more activated in Familiar compared to Unfamiliar tracks. Previous research also found SMA activation during listening to Familiar tracks with the interpretation that participants might be mentally singing along to or engaging more with the Familiar tracks (Freitas et al., 2018; Halpern & Zatorre, 1999; Pereira et al., 2011) or feeling the urge to move to music (Engel et al., 2022). Frontal areas were also more activated by Familiar music including the posterior dorsal cingulate gyrus suggesting the recollection of prior memories (Addis et al., 2007; Rolls, 2019) and the amygdala, pregenual cingulate gyrus and parahippocampal gyrus possibly relating to emotion processing (Palomero-Gallagher & Amunts, 2022). It is logical that listening to Familiar would evoke more memories or use memory-related areas more than Unfamiliar music, these memories are sometimes linked to emotions through episodic memories. Lyrics might be processed faster for Familiar music indicating that the meaning of the lyrics affects the listener’s emotions more or that they can feel more emotions for the singer. Finally, there was less activation in the angular gyrus and the middle frontal gyrus (MFG) for Familiar tracks compared to Unfamiliar tracks suggesting less voluntary attention was required to process these songs, as they were well-known (Li & Tsai, 2024; Rossi et al., 2014). If a song is well known by a listener, that listener does not need to generate as many predictions and therefore can pay less attention to the individual stimuli (Koelsch et al., 2019; Li & Tsai, 2024; Vuust et al., 2022). This could explain why we find less activation in attention areas for Familiar music.

### Music for Relaxation: Individual differences

While there was a stable effect of Familiarity and Type across all participants, some participants had different patterns of relaxation depending on the music. The clustering analysis revealed 4 distinct groups of participants that responded differently to the effect of music Familiarity and Type. One group rated more relaxation for Familiar tracks regardless of Type (Familiarity-driven). One group rated more relaxation for Sedative tracks, regardless of Familiarity (Type-driven). One group rated relaxation the same as the overall results, rating more relaxation for Familiar and Sedative music (Familiarity and Type driven). The last group had inconsistent effects, rating more relaxation for Familiar music but slightly less for Sedative music which is opposite to all other cases (Unclear-drive).

These differences were also apparent in the neuroimaging results. When listening to Excitative compared to Sedative music, Type-driven individuals had more activity in Heschl’s gyrus whereas the other clusters had more peripheral auditory activations. This suggests that these individuals needed to process the auditory stimuli more than the others. This could be due to different processing strategies in different individuals or to differences in embodiment of stimuli (Farina, 2021). Furthermore, when listening to Familiar compared to Unfamiliar music all clusters had activation in motor areas, such as the pre-SMA, Precentral Gyrus and Cerebellum except Type-driven individuals. As with the overall results, these motor activations could be linked to feeling the urge to move to music, or singing along covertly (Freitas et al., 2018; Halpern & Zatorre, 1999; Matthews et al., 2020; Pereira et al., 2011; Scarratt et al., in preparation; Zatorre et al., 2007). This further indicates that Type-driven individuals were listening to the music in a different way, engaging less motor activity.

Type-driven individuals did not find the Familiar music more relaxing, suggesting that motor involvement for Familiar music increases relaxation, or that if individuals are more relaxed, they feel more of an urge to move to sing along to the music (Bella et al., 2015). For example, singing is used in therapy to increase relaxation and mood which we could be seeing in effect here (Bella et al., 2015; Bernardi et al., 2017; Countinho et al., 2014; Kang et al., 2018). The Type-driven cluster also had more activity in the Operculum 9 and 10 for Familiar tracks, areas associated with speech motor planning (Mălîia et al., 2018; Unger et al., 2023). In contrast, clusters with Unclear-driven and Familiarity-driven had more activity for Familiar music in the aSTG and IFG, respectively. This would indicate that the Type-driven individuals process the lyrics of Familiar music differently than the others. Finally, Familiarity-driven individuals had more activation of the MFG when listening to unfamiliar music, suggesting increased attention to the Unfamiliar tracks. This would explain why participants in this cluster was not relaxed by Unfamiliar music as they paid more attention to it.

Thus, our results show key differences in the processing of the auditory stimuli which relates to the behavioural results. Some individuals (Type-driven) process music at lower auditory levels, which might explain why they do not feel relaxed by Excitative, more musically intense music. These same individuals may process lyrics differently and pay as much attention to Familiar as to Unfamiliar tracks, relating to how they do not find comfort in familiarity. Understanding that the same music can affect individuals in different ways is of great importance when deciding which music to recommend or use for relaxation or relaxation-related activities. Music perception can be influenced by listener attributes such as music empathy, absorption (Garrido & Schubert, 2011), music reward (Mas-Herrero et al., 2013), alexithymia (Taruffi et al., 2017) and personality traits (Ladning & Schellenberg, 2012) or situational factors such as physical factors like location and acoustics, or social factors like listening together or alone (Gabrielsson, 2001). Many of these cannot be controlled for in one single experiment. Nevertheless, moving from aggregating all participants together to considering inter-individual differences will lead to a better understanding of listening habits and ultimately more effective recommendation systems for many real-life situations including relaxation.

### Limitations

This study uses music excerpts to investigate how different types of music affects individuals’ states. To be able to have many trials and different tracks in the MRI scanner, the music was cut to 28s. This is not representative of how individuals actually use music. It is possible that some people would need longer than this amount of time to really feel relaxed. However, as we find an effect even in the short excerpts, we also expect to find this effect using longer music tracks. Another limitation is that this study was carried out in the MRI scanner in order to collect brain correlates. MRI machines produce a lot of noise which could have influenced how these music excepts were perceived. However, the noise was the same for all participants and all music conditions and thereby does not explain the effects of the conditions. Finally, it is worth noting that the participants in this study were not purposefully subjected to a stressor. It is possible that a stronger effect could have been found if all participants had undergone a similar stressful task. Given that we still found an effect of music on relaxation, we can conclude that music influences relaxation regardless of baseline stress.

## Conclusion

This study shows how listening to different types of music influence relaxation levels. Globally, listening to sedative and familiar music led to more relaxation. Excitative music required more auditory processing, and unfamiliar music required more attention, explaining why these types of music were found to be the least relaxing. Interestingly, we show behavioural differences between participants in their relaxation responses to the music, which was also reflected in brain activity. This exemplifies how different listener attributes or situational factors influence music perception in the context of relaxation. This is the first study finding individual differences in neural activity in response to music-based relaxation, an exciting way to gain insights in how music is processed. We show strong evidence for considering the interplay between music type, familiarity, and listening styles for music interventions and music neuroimaging studies.

## Supporting information

All supplementary material

