## Supplementary material for "Individual differences in the effects of musical familiarity and musical features on brain activity during relaxation": All supplementary material

| Supplementary Table S1: Characteristics of participants | | |
| --- | --- | --- |
| Nationality | Danish: 11  Eastern Europe: 18 | Western Europe: 19  Non-Western: 9 |
| Current country of residence | Denmark: 51  Hungary: 1  Lithuania: 1 | Poland: 3  Slovenia: 1  The Netherlands: 1 |
| Mean age (SD) | 25.55(4.61) |  |
| Gender | Female: 40  Male: 17 | Non-binary: 1 |
| Handedness | Left: 2  Right: 52 | Ambidextrous: 4 |
| Education | High school: 17  Bachelor: 22  More: 19 |  |
| Singing ability | Poor: 10  Medium: 20 | Group: 16  Good: 12 |
| Frequency music relax | Rarely: 1  Sometimes: 10 | Often: 25  Always: 22 |
| Mean extraversion (SD) | 5.98(2.89) |  |
| Mean Music Training | 2.5(1.60) |  |
| Music genres | Indie: 33  R&B: 11  Lo-Fi: 14  House: 8  Meditation: 10  Pop: 41  EDM: 8  Rap: 11  Country: 7  Reggae: 5 | Folk: 9  Jazz: 10  New Age: 5  Soundtrack: 11  Rock: 15  Punk: 4  World: 5  Gospel: 1  Funk: 4 |
| Type music relax | Neo-soul: 1  Lofi: 5  Indie: 7  Psychedelic: 2  Instrumental: 6  Acoustic: 2  Alternative Rock: 1  Binaural Beats: 1  Classic Rock: 2  Opera: 1  Classical Music: 3  Meditation Music: 2  R&B: 1  Pop: 6  Country: 2  Jazz: 5 | Chillwave: 1  Electronic Music: 2  Folk: 3  IDM: 1  Rock: 3  Blues: 1  Sad or Slow Music: 1  Soft Pop: 1  Soft Rock: 2  Soul: 1  Soundtrack: 2  Trance: 1  Metal: 1  Oriental: 1  Kpop: 1  Jpop: 1  Rap: 2 |

### Supplementary Table S2: Questionnaire

1. I engaged in regular, daily practice of a musical instrument (including voice) for ___ years.	0;1;2;3;4-5;6-9;10 or more
2. At the peak of my interest, I practiced ___ hours per day on my primary instrument.	0;0.5;1;1.5;2;3-4;5 or more
3. I have had formal training in music theory for __ years	0;0.5;1;2;3;4-6;7 or more
4. I have had __ years of formal training on a musical instrument (including voice) during my lifetime.	0;0.5;1;2;3-5;6-9;10 or more
5. I can play ___ musical instruments	0;1;2;3;4;5;6 or more
6. How would you classify yourself as a singer?
   a. Hearing me sing is not a good idea
   b. I’m only good at singing in the shower
   c. I get by in a group
   d. Amateur singer
   e. Professional singer
7. What genres of music do you usually listen to? (select 5 at max)

Country, EDM, Folk, Funk, Gospel, House, Jazz, Indie, Lo-Fi, Meditation, New age, Opera, Pop, Punk, Rap, Reggae, Rock, R&B, Soundtrack, World

8. How often do you use music to relax?

Never, Rarely, Sometimes, Often, Always

9. What type of music do you usually find relaxing?

____

10. Extraversion sub-scale of Eysenck personality test
11. What is your highest level of completed education?

High school, bachelor, master, PhD, More

12. In what year were you born?

____

13. What gender do you identify with?

Male, Female, Non-binary, Other

14. In what country do you live?
15. In what country/countries did you spend most of your childhood (until 18years)?
16. What is your nationality?
17. Are you right-handed?

Yes/No/Ambidextrous

### Estimates for each bayesian model

#### Supplementary Table S3: Basic Bayesian Model

| effect | component | group | term | estimate | std.error | conf.low | conf.high |
| --- | --- | --- | --- | --- | --- | --- | --- |
| fixed | cond | NA | (Intercept) | 0.552797 | 0.086469 | 0.382391 | 0.721665 |
| fixed | cond | NA | phi_(Intercept) | 1.234501 | 0.131784 | 0.971386 | 1.489198 |
| fixed | cond | NA | TypeSedative | 0.232464 | 0.028579 | 0.176869 | 0.288977 |
| fixed | cond | NA | new_familiarityunfamiliar | -0.48657 | 0.028392 | -0.54257 | -0.43102 |
| fixed | cond | NA | phi_TypeSedative | 0.082881 | 0.054389 | -0.02322 | 0.186036 |
| fixed | cond | NA | phi_new_familiarityunfamiliar | 0.453739 | 0.051772 | 0.350054 | 0.554001 |
| ran_pars | cond | fmri_id | sd__(Intercept) | 0.621503 | 0.065348 | 0.509886 | 0.760849 |
| ran_pars | cond | fmri_id | sd__phi_(Intercept) | 0.939837 | 0.094541 | 0.775501 | 1.13333 |

#### Supplementary Table S4: Basic Bayesian Model and Random Effects

| effect | component | group | term | estimate | std.error | conf.low | conf.high |
| --- | --- | --- | --- | --- | --- | --- | --- |
| fixed | cond | NA | (Intercept) | 0.534061 | 0.116694 | 0.306482 | 0.766738 |
| fixed | cond | NA | phi_(Intercept) | 1.469994 | 0.13226 | 1.211344 | 1.733285 |
| fixed | cond | NA | TypeSedative | 0.393858 | 0.079622 | 0.23835 | 0.55651 |
| fixed | cond | NA | new_familiarityunfamiliar | -0.54572 | 0.071817 | -0.68583 | -0.40863 |
| fixed | cond | NA | phi_TypeSedative | 0.126914 | 0.078593 | -0.02528 | 0.28182 |
| fixed | cond | NA | phi_new_familiarityunfamiliar | 0.349195 | 0.074073 | 0.200685 | 0.490996 |
| ran_pars | cond | fmri_id | sd__(Intercept) | 0.84645 | 0.089474 | 0.688863 | 1.040913 |
| ran_pars | cond | fmri_id | sd__TypeSedative | 0.552921 | 0.067706 | 0.441897 | 0.705831 |
| ran_pars | cond | fmri_id | sd__new_familiarityunfamiliar | 0.459499 | 0.06263 | 0.348255 | 0.591051 |
| ran_pars | cond | fmri_id | sd__phi_(Intercept) | 0.973934 | 0.107213 | 0.781236 | 1.205048 |
| ran_pars | cond | fmri_id | sd__phi_TypeSedative | 0.402579 | 0.09095 | 0.226386 | 0.582476 |
| ran_pars | cond | fmri_id | sd__phi_new_familiarityunfamiliar | 0.361346 | 0.098054 | 0.131831 | 0.545734 |
| ran_pars | cond | fmri_id | cor__(Intercept).TypeSedative | -0.31302 | 0.138159 | -0.55801 | -0.03254 |
| ran_pars | cond | fmri_id | cor__(Intercept).new_familiarityunfamiliar | -0.66157 | 0.100541 | -0.82008 | -0.43494 |
| ran_pars | cond | fmri_id | cor__TypeSedative.new_familiarityunfamiliar | 0.13285 | 0.161652 | -0.18881 | 0.441027 |
| ran_pars | cond | fmri_id | cor__phi_(Intercept).phi_TypeSedative | 0.08117 | 0.204596 | -0.30071 | 0.49179 |
| ran_pars | cond | fmri_id | cor__phi_(Intercept).phi_new_familiarityunfamiliar | -0.28275 | 0.206321 | -0.64645 | 0.145077 |
| ran_pars | cond | fmri_id | cor__phi_TypeSedative.phi_new_familiarityunfamiliar | -0.30497 | 0.266332 | -0.78155 | 0.258537 |

#### Supplementary Table S5: Basic Bayesian Model with Liking

| effect | component | group | term | estimate | std.error | conf.low | conf.high |
| --- | --- | --- | --- | --- | --- | --- | --- |
| fixed | cond | NA | (Intercept) | 0.410405 | 0.084328 | 0.243535 | 0.575495 |
| fixed | cond | NA | phi_(Intercept) | 1.351486 | 0.124109 | 1.101839 | 1.588861 |
| fixed | cond | NA | TypeSedative | 0.263805 | 0.029456 | 0.205944 | 0.323044 |
| fixed | cond | NA | new_familiarityunfamiliar | -0.21895 | 0.034393 | -0.28813 | -0.15106 |
| fixed | cond | NA | Liking_z | 0.289624 | 0.019898 | 0.251122 | 0.327994 |
| fixed | cond | NA | phi_TypeSedative | 0.102564 | 0.054765 | -0.00343 | 0.210702 |
| fixed | cond | NA | phi_new_familiarityunfamiliar | 0.360795 | 0.062341 | 0.237347 | 0.482457 |
| fixed | cond | NA | phi_Liking_z | -0.03618 | 0.030383 | -0.09246 | 0.024418 |
| ran_pars | cond | fmri_id | sd__(Intercept) | 0.574807 | 0.062717 | 0.47022 | 0.713194 |
| ran_pars | cond | fmri_id | sd__phi_(Intercept) | 0.914299 | 0.090125 | 0.760278 | 1.109744 |

#### Supplementary Table S6: Basic Bayesian Model with Liking and Random Effects

| effect | component | group | term | estimate | std.error | conf.low | conf.high |
| --- | --- | --- | --- | --- | --- | --- | --- |
| fixed | cond | NA | (Intercept) | 0.354285 | 0.110549 | 0.137305 | 0.572199 |
| fixed | cond | NA | phi_(Intercept) | 1.566096 | 0.137715 | 1.289492 | 1.842857 |
| fixed | cond | NA | TypeSedative | 0.452687 | 0.081771 | 0.288362 | 0.612892 |
| fixed | cond | NA | new_familiarityunfamiliar | -0.24058 | 0.070455 | -0.38191 | -0.10627 |
| fixed | cond | NA | Liking_z | 0.317373 | 0.019915 | 0.279256 | 0.356514 |
| fixed | cond | NA | phi_TypeSedative | 0.189826 | 0.085196 | 0.024369 | 0.361402 |
| fixed | cond | NA | phi_new_familiarityunfamiliar | 0.263056 | 0.076046 | 0.108631 | 0.413767 |
| fixed | cond | NA | phi_Liking_z | -0.00429 | 0.033387 | -0.06986 | 0.060516 |
| ran_pars | cond | fmri_id | sd__(Intercept) | 0.824061 | 0.083879 | 0.675867 | 1.004277 |
| ran_pars | cond | fmri_id | sd__TypeSedative | 0.560753 | 0.065391 | 0.448673 | 0.706567 |
| ran_pars | cond | fmri_id | sd__new_familiarityunfamiliar | 0.466355 | 0.06088 | 0.36069 | 0.594985 |
| ran_pars | cond | fmri_id | sd__phi_(Intercept) | 0.975807 | 0.109164 | 0.788167 | 1.205508 |
| ran_pars | cond | fmri_id | sd__phi_TypeSedative | 0.476254 | 0.084916 | 0.315162 | 0.649168 |
| ran_pars | cond | fmri_id | sd__phi_new_familiarityunfamiliar | 0.264582 | 0.109466 | 0.035825 | 0.459822 |
| ran_pars | cond | fmri_id | cor__(Intercept).TypeSedative | -0.34091 | 0.132046 | -0.58099 | -0.06803 |
| ran_pars | cond | fmri_id | cor__(Intercept).new_familiarityunfamiliar | -0.74265 | 0.079359 | -0.86975 | -0.55825 |
| ran_pars | cond | fmri_id | cor__TypeSedative.new_familiarityunfamiliar | 0.18827 | 0.156408 | -0.12246 | 0.481483 |
| ran_pars | cond | fmri_id | cor__phi_(Intercept).phi_TypeSedative | -0.11492 | 0.179719 | -0.45674 | 0.243927 |
| ran_pars | cond | fmri_id | cor__phi_(Intercept).phi_new_familiarityunfamiliar | -0.25203 | 0.26044 | -0.70758 | 0.345416 |
| ran_pars | cond | fmri_id | cor__phi_TypeSedative.phi_new_familiarityunfamiliar | -0.25594 | 0.312093 | -0.81309 | 0.39256 |

#### Supplementary Table S7: Basic Bayesian Model with Liking and Random Effects and Singing Ability

| effect | component | group | term | estimate | std.error | conf.low | conf.high |
| --- | --- | --- | --- | --- | --- | --- | --- |
| fixed | cond | NA | (Intercept) | 0.844415 | 0.180965 | 0.4838 | 1.189334 |
| fixed | cond | NA | phi_(Intercept) | 1.544577 | 0.319683 | 0.913132 | 2.183704 |
| fixed | cond | NA | TypeSedative | 0.44945 | 0.080908 | 0.28994 | 0.611504 |
| fixed | cond | NA | new_familiarityunfamiliar | -0.24099 | 0.075225 | -0.38933 | -0.09123 |
| fixed | cond | NA | Liking_z | 0.31677 | 0.020427 | 0.277481 | 0.356721 |
| fixed | cond | NA | singing_abilityGroup | -0.51918 | 0.203218 | -0.91348 | -0.10534 |
| fixed | cond | NA | singing_abilityGood | -0.57698 | 0.215264 | -0.99317 | -0.13342 |
| fixed | cond | NA | singing_abilityMedium | -0.65459 | 0.20299 | -1.02716 | -0.22495 |
| fixed | cond | NA | phi_TypeSedative | 0.187515 | 0.086115 | 0.016763 | 0.35669 |
| fixed | cond | NA | phi_new_familiarityunfamiliar | 0.268875 | 0.07884 | 0.113132 | 0.422899 |
| fixed | cond | NA | phi_Liking_z | -0.00521 | 0.033739 | -0.07136 | 0.060916 |
| fixed | cond | NA | phi_singing_abilityGroup | 0.359823 | 0.391717 | -0.4027 | 1.123482 |
| fixed | cond | NA | phi_singing_abilityGood | -0.09835 | 0.420282 | -0.95952 | 0.698683 |
| fixed | cond | NA | phi_singing_abilityMedium | -0.17164 | 0.387631 | -0.95983 | 0.575938 |
| ran_pars | cond | fmri_id | sd__(Intercept) | 0.748001 | 0.083945 | 0.603766 | 0.936201 |
| ran_pars | cond | fmri_id | sd__TypeSedative | 0.565316 | 0.066681 | 0.444642 | 0.706747 |
| ran_pars | cond | fmri_id | sd__new_familiarityunfamiliar | 0.466801 | 0.061884 | 0.355875 | 0.597151 |
| ran_pars | cond | fmri_id | sd__phi_(Intercept) | 0.965241 | 0.108137 | 0.776245 | 1.197088 |
| ran_pars | cond | fmri_id | sd__phi_TypeSedative | 0.474833 | 0.084335 | 0.313791 | 0.644211 |
| ran_pars | cond | fmri_id | sd__phi_new_familiarityunfamiliar | 0.277276 | 0.104632 | 0.047882 | 0.469371 |
| ran_pars | cond | fmri_id | cor__(Intercept).TypeSedative | -0.34944 | 0.133161 | -0.58371 | -0.06821 |
| ran_pars | cond | fmri_id | cor__(Intercept).new_familiarityunfamiliar | -0.73811 | 0.083132 | -0.8715 | -0.54634 |
| ran_pars | cond | fmri_id | cor__TypeSedative.new_familiarityunfamiliar | 0.196547 | 0.15542 | -0.1107 | 0.494159 |
| ran_pars | cond | fmri_id | cor__phi_(Intercept).phi_TypeSedative | -0.03553 | 0.191882 | -0.40221 | 0.347008 |
| ran_pars | cond | fmri_id | cor__phi_(Intercept).phi_new_familiarityunfamiliar | -0.24565 | 0.250099 | -0.68592 | 0.323419 |
| ran_pars | cond | fmri_id | cor__phi_TypeSedative.phi_new_familiarityunfamiliar | -0.27089 | 0.302369 | -0.8013 | 0.382839 |

#### Supplementary Table S8: Zero inflated Bayesian Model

| effect | component | group | term | estimate | std.error | conf.low | conf.high |
| --- | --- | --- | --- | --- | --- | --- | --- |
| fixed | cond | NA | (Intercept) | 0.491648 | 0.077641 | 0.33972 | 0.644852 |
| fixed | cond | NA | phi_(Intercept) | 1.597149 | 0.114144 | 1.372956 | 1.825469 |
| fixed | cond | NA | zoi_(Intercept) | -3.29276 | 0.33974 | -3.9959 | -2.6595 |
| fixed | cond | NA | coi_(Intercept) | 2.593102 | 0.928026 | 0.997407 | 4.592884 |
| fixed | cond | NA | TypeSedative | 0.236697 | 0.028483 | 0.181792 | 0.293352 |
| fixed | cond | NA | new_familiarityunfamiliar | -0.43935 | 0.027591 | -0.49281 | -0.38617 |
| fixed | cond | NA | phi_TypeSedative | 0.15503 | 0.059702 | 0.038419 | 0.273503 |
| fixed | cond | NA | phi_new_familiarityunfamiliar | 0.258348 | 0.054956 | 0.152299 | 0.366663 |
| fixed | cond | NA | zoi_TypeSedative | 0.250317 | 0.171728 | -0.08067 | 0.585145 |
| fixed | cond | NA | zoi_new_familiarityunfamiliar | -1.68532 | 0.210815 | -2.09706 | -1.28722 |
| fixed | cond | NA | coi_TypeSedative | 1.722013 | 0.790657 | 0.302085 | 3.442805 |
| fixed | cond | NA | coi_new_familiarityunfamiliar | -3.73884 | 0.994031 | -5.85196 | -1.93941 |
| ran_pars | cond | fmri_id | sd__(Intercept) | 0.525841 | 0.056386 | 0.427894 | 0.647705 |
| ran_pars | cond | fmri_id | sd__phi_(Intercept) | 0.755071 | 0.079517 | 0.614539 | 0.924356 |
| ran_pars | cond | fmri_id | sd__zoi_(Intercept) | 1.883235 | 0.317512 | 1.362798 | 2.586309 |
| ran_pars | cond | fmri_id | sd__coi_(Intercept) | 4.020867 | 1.189622 | 2.185917 | 6.68924 |

#### Supplementary Table S9: Zero inflated Bayesian Model with Liking

| effect | component | group | term | estimate | std.error | conf.low | conf.high |
| --- | --- | --- | --- | --- | --- | --- | --- |
| fixed | cond | NA | (Intercept) | 0.413723 | 0.083964 | 0.247837 | 0.572414 |
| fixed | cond | NA | phi_(Intercept) | 1.337754 | 0.137375 | 1.066403 | 1.610645 |
| fixed | cond | NA | zoi_(Intercept) | -10.9969 | 3.097224 | -18.378 | -6.67042 |
| fixed | cond | NA | coi_(Intercept) | 21292261 | 89259262 | -8.2E+07 | 2.28E+08 |
| fixed | cond | NA | TypeSedative | 0.26394 | 0.028966 | 0.208689 | 0.320562 |
| fixed | cond | NA | new_familiarityunfamiliar | -0.21949 | 0.033299 | -0.28334 | -0.15754 |
| fixed | cond | NA | Liking_z | 0.28881 | 0.019487 | 0.251022 | 0.328053 |
| fixed | cond | NA | phi_TypeSedative | 0.102079 | 0.055426 | -0.00644 | 0.208821 |
| fixed | cond | NA | phi_new_familiarityunfamiliar | 0.360036 | 0.061124 | 0.244867 | 0.480659 |
| fixed | cond | NA | phi_Liking_z | -0.03637 | 0.030019 | -0.09478 | 0.023797 |
| fixed | cond | NA | zoi_TypeSedative | 0.02262 | 3.107105 | -6.37806 | 6.433251 |
| fixed | cond | NA | zoi_new_familiarityunfamiliar | 0.246212 | 3.363175 | -6.48712 | 6.958169 |
| fixed | cond | NA | zoi_Liking_z | 0.300478 | 1.317823 | -2.02084 | 3.206077 |
| fixed | cond | NA | coi_TypeSedative | 42163557 | 60709299 | -2.8E+07 | 1.9E+08 |
| fixed | cond | NA | coi_new_familiarityunfamiliar | -8.5E+07 | 1.61E+08 | -4.9E+08 | 23348365 |
| fixed | cond | NA | coi_Liking_z | 4469578 | 54966991 | -1.1E+08 | 1.35E+08 |
| ran_pars | cond | fmri_id | sd__(Intercept) | 0.569526 | 0.05687 | 0.469661 | 0.693724 |
| ran_pars | cond | fmri_id | sd__phi_(Intercept) | 0.929944 | 0.096327 | 0.768152 | 1.144272 |
| ran_pars | cond | fmri_id | sd__zoi_(Intercept) | 0.79404 | 0.631496 | 0.030437 | 2.375641 |
| ran_pars | cond | fmri_id | sd__coi_(Intercept) | 2.757281 | 3.229545 | 0.074038 | 10.34728 |

#### Supplementary Table S10: Zero inflated Bayesian Model with Liking and Random Effects

| effect | component | group | term | estimate | std.error | conf.low | conf.high |
| --- | --- | --- | --- | --- | --- | --- | --- |
| fixed | cond | NA | (Intercept) | 0.359453 | 0.117344 | 0.138863 | 0.600133 |
| fixed | cond | NA | phi_(Intercept) | 1.570326 | 0.14317 | 1.274463 | 1.848043 |
| fixed | cond | NA | zoi_(Intercept) | -11.173 | 3.248948 | -19.332 | -6.60616 |
| fixed | cond | NA | coi_(Intercept) | 3525681 | 8150831 | -2024750 | 26482445 |
| fixed | cond | NA | TypeSedative | 0.444629 | 0.08179 | 0.287243 | 0.602215 |
| fixed | cond | NA | new_familiarityunfamiliar | -0.24162 | 0.074663 | -0.39412 | -0.09996 |
| fixed | cond | NA | Liking_z | 0.317303 | 0.019935 | 0.27826 | 0.356303 |
| fixed | cond | NA | phi_TypeSedative | 0.180345 | 0.087067 | 0.007505 | 0.352352 |
| fixed | cond | NA | phi_new_familiarityunfamiliar | 0.267365 | 0.079933 | 0.109073 | 0.423733 |
| fixed | cond | NA | phi_Liking_z | -0.00378 | 0.03313 | -0.06991 | 0.060078 |
| fixed | cond | NA | zoi_TypeSedative | -0.65096 | 3.519883 | -8.03905 | 6.288318 |
| fixed | cond | NA | zoi_new_familiarityunfamiliar | -0.34248 | 3.471501 | -7.20065 | 6.666976 |
| fixed | cond | NA | zoi_Liking_z | 0.260638 | 1.373541 | -2.15476 | 3.31689 |
| fixed | cond | NA | coi_TypeSedative | -5196949 | 14974985 | -5E+07 | 4671916 |
| fixed | cond | NA | coi_new_familiarityunfamiliar | -1865419 | 5595923 | -2E+07 | 5138932 |
| fixed | cond | NA | coi_Liking_z | -505025 | 941824.3 | -2337742 | 1062549 |
| ran_pars | cond | fmri_id | sd__(Intercept) | 0.821626 | 0.083721 | 0.675084 | 1.002582 |
| ran_pars | cond | fmri_id | sd__TypeSedative | 0.565196 | 0.06774 | 0.445668 | 0.709472 |
| ran_pars | cond | fmri_id | sd__new_familiarityunfamiliar | 0.464701 | 0.060576 | 0.354495 | 0.59107 |
| ran_pars | cond | fmri_id | sd__phi_(Intercept) | 0.971526 | 0.106563 | 0.786121 | 1.200498 |
| ran_pars | cond | fmri_id | sd__phi_TypeSedative | 0.471804 | 0.084154 | 0.309532 | 0.642152 |
| ran_pars | cond | fmri_id | sd__phi_new_familiarityunfamiliar | 0.280736 | 0.105716 | 0.049335 | 0.474534 |
| ran_pars | cond | fmri_id | sd__zoi_(Intercept) | 0.917449 | 0.728376 | 0.032296 | 2.742879 |
| ran_pars | cond | fmri_id | sd__zoi_TypeSedative | 1.162283 | 0.957602 | 0.033476 | 3.611642 |
| ran_pars | cond | fmri_id | sd__zoi_new_familiarityunfamiliar | 1.151134 | 0.962075 | 0.049985 | 3.580362 |
| ran_pars | cond | fmri_id | sd__coi_(Intercept) | 2.711486 | 3.000216 | 0.098606 | 10.17785 |
| ran_pars | cond | fmri_id | sd__coi_TypeSedative | 2.707094 | 3.021147 | 0.105034 | 9.803726 |
| ran_pars | cond | fmri_id | sd__coi_new_familiarityunfamiliar | 2.733989 | 3.362028 | 0.078278 | 10.14938 |
| ran_pars | cond | fmri_id | cor__(Intercept).TypeSedative | -0.34841 | 0.132927 | -0.58826 | -0.08154 |
| ran_pars | cond | fmri_id | cor__(Intercept).new_familiarityunfamiliar | -0.744 | 0.078806 | -0.86751 | -0.56594 |
| ran_pars | cond | fmri_id | cor__TypeSedative.new_familiarityunfamiliar | 0.202592 | 0.153251 | -0.10775 | 0.487764 |
| ran_pars | cond | fmri_id | cor__phi_(Intercept).phi_TypeSedative | -0.10255 | 0.180545 | -0.43736 | 0.265151 |
| ran_pars | cond | fmri_id | cor__phi_(Intercept).phi_new_familiarityunfamiliar | -0.26385 | 0.243954 | -0.70006 | 0.282961 |
| ran_pars | cond | fmri_id | cor__phi_TypeSedative.phi_new_familiarityunfamiliar | -0.25204 | 0.310295 | -0.81536 | 0.399558 |
| ran_pars | cond | fmri_id | cor__zoi_(Intercept).zoi_TypeSedative | -0.0716 | 0.504008 | -0.90205 | 0.846961 |
| ran_pars | cond | fmri_id | cor__zoi_(Intercept).zoi_new_familiarityunfamiliar | -0.07753 | 0.50462 | -0.91296 | 0.852855 |
| ran_pars | cond | fmri_id | cor__zoi_TypeSedative.zoi_new_familiarityunfamiliar | -0.04308 | 0.49336 | -0.8832 | 0.860867 |
| ran_pars | cond | fmri_id | cor__coi_(Intercept).coi_TypeSedative | 0.00356 | 0.494864 | -0.8815 | 0.884288 |
| ran_pars | cond | fmri_id | cor__coi_(Intercept).coi_new_familiarityunfamiliar | -0.00364 | 0.501382 | -0.87345 | 0.874817 |
| ran_pars | cond | fmri_id | cor__coi_TypeSedative.coi_new_familiarityunfamiliar | 0.012344 | 0.503504 | -0.88075 | 0.87727 |

#### Supplementary Table S11: Comparison between models based on P_WAIC, wAIC and ELPD_WAIC

| Model | P_waic  Estimate(sd) | Waic Estimate(sd) | Elpd_waic |
| --- | --- | --- | --- |
| Fancy_model_z | 167.9(8.6) | -834.4(97.9) | 417.2(48.9) |
| Fancy_model_liking | 128.1(7.3) | -2658.4(95.3) | 1329.2(47.7) |
| Fancy_model_liking_id | 262.3(12.0) | -3061.3(99.2) | 1530.7(49.6) |
| Simple | 127.0(7.4) | -2442.3(93.8) | 1221.2(46.9) |
| Simple_id | 261.7(12.0) | -2813.6(98.5) | 1406.8(49.3) |
| Model_beta_simple_liking | 129.3(7.5) | -2658.6(95.4) | 1329.3(47.7) |
| Model_beta_simple_liking_id | 261.1(12.1) | -3064.1(99.3) | 1532.0(49.7) |
| Model_beta_simple_liking_id_singing | 261.3(12.0) | -3065.8(99.4) | 1532.0(49.7) |

### GLM beta regression

#### Supplementary Analysis S1

We performed the same analysis using linear regression with a beta distribution using the R package GlmmTMB with stepwise models using the same variables as for the Zero-One Inflated Bayesian analysis and adding confounders. The details of all the models are in Supplementary Table XX and the AIC and BIC values for the significant models are in Supplementary Table XX. The model that performed best was Relaxation ~ Type + Familiarity + Liking + Singing Ability + (1+Familiarity+Type|fmri_id).

The general linear beta regression model reveals that the intercept of Relaxation for Excitative Familiar music is equal to **55.8%** (z = 1.586), which increases by **11.2%** (z = 5.678, p<.001) if the music is Sedative, and decreases by **4.6%** (z = -2.291, p<.05) if the music is Unfamiliar. Additionally, Relaxation increases by **9.6%** (z = 17.571, p<.001) with each unit increase in Liking. The model also found a significant effect for singing ability. Compared to individuals who said they could sing well in a group, the Relaxation levels of individuals who indicated that they could not sing increased by **13.4%.** The other singing ability levels were not different from the comparator, indicating that higher singing ability levels correlate with lower relaxation scores. The numbers for the estimates in logit are in Supplementary Table X.

It is important to note that singing ability was only measured by one self-reported question so it should not be given too much importance. The estimates for the model that does not include singing ability has very similar results. The intercept of Relaxation for Excitative Familiar music is equal to **57.7%** (z = 2.667, p = 0.00766), which increases by **11%** (z = 5.681, p < .001) if the music is Sedative, and decreases by **4.6%** (z = -2.279, p < .05) if the music is Unfamiliar. Additionally, Relaxation increases by **9.4%** (z = 17.555, p < .001) with each unit increase in Liking. Thus, the results are robust to the inclusion or exclusion of singing ability. Furthermore, the results are also robust to the modelling method as both the GLM and Bayesian analysis reveal similar effects of Type, Familiarity and Liking.

#### Estimates for all GLM models

##### Supplementary Table S12: **Type + Familiarity**

| effect | component | group | term | estimate | std.error | statistic | p.value |
| --- | --- | --- | --- | --- | --- | --- | --- |
| fixed | cond | NA | (Intercept) | 0.533554 | 0.084276 | 6.331048 | 2.44E-10 |
| fixed | cond | NA | TypeSedative | 0.388432 | 0.037892 | 10.25113 | 1.17E-24 |
| fixed | cond | NA | new_familiarityunfamiliar | -0.58104 | 0.038279 | -15.1794 | 4.84E-52 |
| ran_pars | cond | fmri_id | sd__(Intercept) | 0.58903 | NA | NA | NA |

##### Supplementary Table S13: **Type, Familiarity and Random effects**

| effect | component | group | term | estimate | std.error | statistic | p.value |
| --- | --- | --- | --- | --- | --- | --- | --- |
| fixed | cond | NA | (Intercept) | 0.555802 | 0.117511 | 4.729774 | 2.25E-06 |
| fixed | cond | NA | TypeSedative | 0.394975 | 0.084107 | 4.696076 | 2.65E-06 |
| fixed | cond | NA | new_familiarityunfamiliar | -0.60118 | 0.078065 | -7.70104 | 1.35E-14 |
| ran_pars | cond | fmri_id | sd__(Intercept) | 0.856031 | NA | NA | NA |
| ran_pars | cond | fmri_id | sd__TypeSedative | 0.573355 | NA | NA | NA |
| ran_pars | cond | fmri_id | sd__new_familiarityunfamiliar | 0.52002 | NA | NA | NA |
| ran_pars | cond | fmri_id | cor__(Intercept).TypeSedative | -0.44 | NA | NA | NA |
| ran_pars | cond | fmri_id | cor__(Intercept).new_familiarityunfamiliar | -0.71274 | NA | NA | NA |
| ran_pars | cond | fmri_id | cor__TypeSedative.new_familiarityunfamiliar | 0.150572 | NA | NA | NA |

##### Supplementary Table S14: **Type, Familiarity, Random Effects and Liking**

| effect | component | group | term | estimate | std.error | statistic | p.value |
| --- | --- | --- | --- | --- | --- | --- | --- |
| fixed | cond | NA | (Intercept) | 0.30994 | 0.116221 | 2.666821 | 0.007657 |
| fixed | cond | NA | TypeSedative | 0.47709 | 0.083984 | 5.680715 | 1.34E-08 |
| fixed | cond | NA | new_familiarityunfamiliar | -0.18438 | 0.080897 | -2.27913 | 0.022659 |
| fixed | cond | NA | Liking_z | 0.405685 | 0.02311 | 17.55474 | 5.47E-69 |
| ran_pars | cond | fmri_id | sd__(Intercept) | 0.841456 | NA | NA | NA |
| ran_pars | cond | fmri_id | sd__TypeSedative | 0.57512 | NA | NA | NA |
| ran_pars | cond | fmri_id | sd__new_familiarityunfamiliar | 0.518594 | NA | NA | NA |
| ran_pars | cond | fmri_id | cor__(Intercept).TypeSedative | -0.50748 | NA | NA | NA |
| ran_pars | cond | fmri_id | cor__(Intercept).new_familiarityunfamiliar | -0.76979 | NA | NA | NA |
| ran_pars | cond | fmri_id | cor__TypeSedative.new_familiarityunfamiliar | 0.286004 | NA | NA | NA |

##### Supplementary Table S15: **Type, Familiarity, Random Effects, Liking and Age**

| effect | component | group | term | estimate | std.error | statistic | p.value |
| --- | --- | --- | --- | --- | --- | --- | --- |
| fixed | cond | NA | (Intercept) | 0.336856 | 0.422037 | 0.798168 | 0.424773 |
| fixed | cond | NA | TypeSedative | 0.477065 | 0.083987 | 5.680207 | 1.35E-08 |
| fixed | cond | NA | new_familiarityunfamiliar | -0.18437 | 0.080899 | -2.27901 | 0.022666 |
| fixed | cond | NA | Liking_z | 0.40566 | 0.023114 | 17.55016 | 5.93E-69 |
| fixed | cond | NA | age | -0.00105 | 0.015877 | -0.06633 | 0.947112 |
| ran_pars | cond | fmri_id | sd__(Intercept) | 0.840134 | NA | NA | NA |
| ran_pars | cond | fmri_id | sd__TypeSedative | 0.575139 | NA | NA | NA |
| ran_pars | cond | fmri_id | sd__new_familiarityunfamiliar | 0.518599 | NA | NA | NA |
| ran_pars | cond | fmri_id | cor__(Intercept).TypeSedative | -0.50805 | NA | NA | NA |
| ran_pars | cond | fmri_id | cor__(Intercept).new_familiarityunfamiliar | -0.76849 | NA | NA | NA |
| ran_pars | cond | fmri_id | cor__TypeSedative.new_familiarityunfamiliar | 0.285925 | NA | NA | NA |

##### Supplementary Table S16: **Type, Familiarity, Random Effects, Liking and Extraversion**

| effect | component | group | term | estimate | std.error | statistic | p.value |
| --- | --- | --- | --- | --- | --- | --- | --- |
| fixed | cond | NA | (Intercept) | 0.38083 | 0.181687 | 2.09608 | 0.036075 |
| fixed | cond | NA | TypeSedative | 0.477018 | 0.083983 | 5.679941 | 1.35E-08 |
| fixed | cond | NA | new_familiarityunfamiliar | -0.1841 | 0.080889 | -2.27593 | 0.02285 |
| fixed | cond | NA | Liking_z | 0.405973 | 0.023119 | 17.56032 | 4.96E-69 |
| fixed | cond | NA | extraversion | -0.01184 | 0.023317 | -0.50784 | 0.611567 |
| ran_pars | cond | fmri_id | sd__(Intercept) | 0.841687 | NA | NA | NA |
| ran_pars | cond | fmri_id | sd__TypeSedative | 0.575111 | NA | NA | NA |
| ran_pars | cond | fmri_id | sd__new_familiarityunfamiliar | 0.518502 | NA | NA | NA |
| ran_pars | cond | fmri_id | cor__(Intercept).TypeSedative | -0.51199 | NA | NA | NA |
| ran_pars | cond | fmri_id | cor__(Intercept).new_familiarityunfamiliar | -0.76891 | NA | NA | NA |
| ran_pars | cond | fmri_id | cor__TypeSedative.new_familiarityunfamiliar | 0.286376 | NA | NA | NA |

##### Supplementary Table S17: **Type, Familiarity, Random Effects, Liking and Pop**

| effect | component | group | term | estimate | std.error | statistic | p.value |
| --- | --- | --- | --- | --- | --- | --- | --- |
| fixed | cond | NA | (Intercept) | 0.084496 | 0.182008 | 0.464242 | 0.642474 |
| fixed | cond | NA | TypeSedative | 0.477186 | 0.084 | 5.680777 | 1.34E-08 |
| fixed | cond | NA | new_familiarityunfamiliar | -0.18466 | 0.080942 | -2.28134 | 0.022528 |
| fixed | cond | NA | Liking_z | 0.405292 | 0.023085 | 17.55624 | 5.33E-69 |
| fixed | cond | NA | pop1 | 0.279754 | 0.172178 | 1.624795 | 0.104206 |
| ran_pars | cond | fmri_id | sd__(Intercept) | 0.854796 | NA | NA | NA |
| ran_pars | cond | fmri_id | sd__TypeSedative | 0.575265 | NA | NA | NA |
| ran_pars | cond | fmri_id | sd__new_familiarityunfamiliar | 0.519116 | NA | NA | NA |
| ran_pars | cond | fmri_id | cor__(Intercept).TypeSedative | -0.50959 | NA | NA | NA |
| ran_pars | cond | fmri_id | cor__(Intercept).new_familiarityunfamiliar | -0.79104 | NA | NA | NA |
| ran_pars | cond | fmri_id | cor__TypeSedative.new_familiarityunfamiliar | 0.285299 | NA | NA | NA |

##### Supplementary Table S18: **Type, Familiarity, Random Effects, Liking and Nationality**

| effect | component | group | term | estimate | std.error | statistic | p.value |
| --- | --- | --- | --- | --- | --- | --- | --- |
| fixed | cond | NA | (Intercept) | 0.334291 | 0.183344 | 1.823296 | 0.068259 |
| fixed | cond | NA | TypeSedative | 0.47718 | 0.083983 | 5.681837 | 1.33E-08 |
| fixed | cond | NA | new_familiarityunfamiliar | -0.18424 | 0.080912 | -2.27708 | 0.022782 |
| fixed | cond | NA | Liking_z | 0.405808 | 0.023102 | 17.56617 | 4.47E-69 |
| fixed | cond | NA | cat_nationalityNon- Western | 0.060086 | 0.230949 | 0.260169 | 0.794734 |
| fixed | cond | NA | cat_nationalityWestern | -0.0522 | 0.180432 | -0.28929 | 0.772358 |
| ran_pars | cond | fmri_id | sd__(Intercept) | 0.851519 | NA | NA | NA |
| ran_pars | cond | fmri_id | sd__TypeSedative | 0.575119 | NA | NA | NA |
| ran_pars | cond | fmri_id | sd__new_familiarityunfamiliar | 0.518749 | NA | NA | NA |
| ran_pars | cond | fmri_id | cor__(Intercept).TypeSedative | -0.50892 | NA | NA | NA |
| ran_pars | cond | fmri_id | cor__(Intercept).new_familiarityunfamiliar | -0.77787 | NA | NA | NA |
| ran_pars | cond | fmri_id | cor__TypeSedative.new_familiarityunfamiliar | 0.285978 | NA | NA | NA |

##### Supplementary Table S19: **Type, Familiarity, Random Effects, Liking and Education**

| effect | component | group | term | estimate | std.error | statistic | p.value |
| --- | --- | --- | --- | --- | --- | --- | --- |
| fixed | cond | NA | (Intercept) | 0.364216 | 0.146544 | 2.485367 | 0.012942 |
| fixed | cond | NA | TypeSedative | 0.477146 | 0.083977 | 5.68183 | 1.33E-08 |
| fixed | cond | NA | new_familiarityunfamiliar | -0.1846 | 0.080909 | -2.2816 | 0.022513 |
| fixed | cond | NA | Liking_z | 0.405681 | 0.023107 | 17.55632 | 5.32E-69 |
| fixed | cond | NA | cat_educationHigh school | -0.09599 | 0.168753 | -0.56882 | 0.569477 |
| fixed | cond | NA | cat_educationHigher | -0.07693 | 0.162381 | -0.47374 | 0.635686 |
| ran_pars | cond | fmri_id | sd__(Intercept) | 0.843085 | NA | NA | NA |
| ran_pars | cond | fmri_id | sd__TypeSedative | 0.575064 | NA | NA | NA |
| ran_pars | cond | fmri_id | sd__new_familiarityunfamiliar | 0.518699 | NA | NA | NA |
| ran_pars | cond | fmri_id | cor__(Intercept).TypeSedative | -0.50578 | NA | NA | NA |
| ran_pars | cond | fmri_id | cor__(Intercept).new_familiarityunfamiliar | -0.77411 | NA | NA | NA |
| ran_pars | cond | fmri_id | cor__TypeSedative.new_familiarityunfamiliar | 0.2866 | NA | NA | NA |

##### Supplementary Table S20: **Type, Familiarity, Random Effects, Liking and Singing Ability**

| effect | component | group | term | estimate | std.error | statistic | p.value |
| --- | --- | --- | --- | --- | --- | --- | --- |
| fixed | cond | NA | (Intercept) | 0.195503 | 0.160194 | 1.220413 | 0.222308 |
| fixed | cond | NA | TypeSedative | 0.476973 | 0.084007 | 5.677793 | 1.36E-08 |
| fixed | cond | NA | new_familiarityunfamiliar | -0.18524 | 0.080859 | -2.29093 | 0.021967 |
| fixed | cond | NA | Liking_z | 0.405693 | 0.023089 | 17.57112 | 4.10E-69 |
| fixed | cond | NA | singing_abilityMedium | -0.01265 | 0.173841 | -0.07274 | 0.942013 |
| fixed | cond | NA | singing_abilityGroup | 0.036896 | 0.177746 | 0.207576 | 0.83556 |
| fixed | cond | NA | singing_abilityPoor | 0.616549 | 0.199681 | 3.087665 | 0.002017 |
| ran_pars | cond | fmri_id | sd__(Intercept) | 0.766128 | NA | NA | NA |
| ran_pars | cond | fmri_id | sd__TypeSedative | 0.575298 | NA | NA | NA |
| ran_pars | cond | fmri_id | sd__new_familiarityunfamiliar | 0.518396 | NA | NA | NA |
| ran_pars | cond | fmri_id | cor__(Intercept).TypeSedative | -0.5262 | NA | NA | NA |
| ran_pars | cond | fmri_id | cor__(Intercept).new_familiarityunfamiliar | -0.77002 | NA | NA | NA |
| ran_pars | cond | fmri_id | cor__TypeSedative.new_familiarityunfamiliar | 0.288305 | NA | NA | NA |

##### Supplementary Table S21: **Type, Familiarity, Random Effects, Liking and Music Training**

| effect | component | group | term | estimate | std.error | statistic | p.value |
| --- | --- | --- | --- | --- | --- | --- | --- |
| fixed | cond | NA | (Intercept) | 0.464062 | 0.156552 | 2.964261 | 0.003034 |
| fixed | cond | NA | TypeSedative | 0.477252 | 0.083948 | 5.685078 | 1.31E-08 |
| fixed | cond | NA | new_familiarityunfamiliar | -0.18577 | 0.080869 | -2.29713 | 0.021611 |
| fixed | cond | NA | Liking_z | 0.405771 | 0.023089 | 17.57442 | 3.87E-69 |
| fixed | cond | NA | meantMT | -0.06215 | 0.042347 | -1.46757 | 0.142221 |
| ran_pars | cond | fmri_id | sd__(Intercept) | 0.840043 | NA | NA | NA |
| ran_pars | cond | fmri_id | sd__TypeSedative | 0.574822 | NA | NA | NA |
| ran_pars | cond | fmri_id | sd__new_familiarityunfamiliar | 0.5184 | NA | NA | NA |
| ran_pars | cond | fmri_id | cor__(Intercept).TypeSedative | -0.50382 | NA | NA | NA |
| ran_pars | cond | fmri_id | cor__(Intercept).new_familiarityunfamiliar | -0.7833 | NA | NA | NA |
| ran_pars | cond | fmri_id | cor__TypeSedative.new_familiarityunfamiliar | 0.289854 | NA | NA | NA |

| Supplementary Table S22: GLM Model Comparison | | |
| --- | --- | --- |
| Model | AIC | BIC |
| 1 Type + Familiarity | -1344.726 | -1315.199 |
| 2 Type, Familiarity + ID | -1602.368 | -1543.314 |
| 3 Type, Familiarity, ID + Liking | -1896.631 | -1831.671 |
| 4 Type, Familiarity, ID, Liking +singing ability | -1903.047 | -1820.370 |

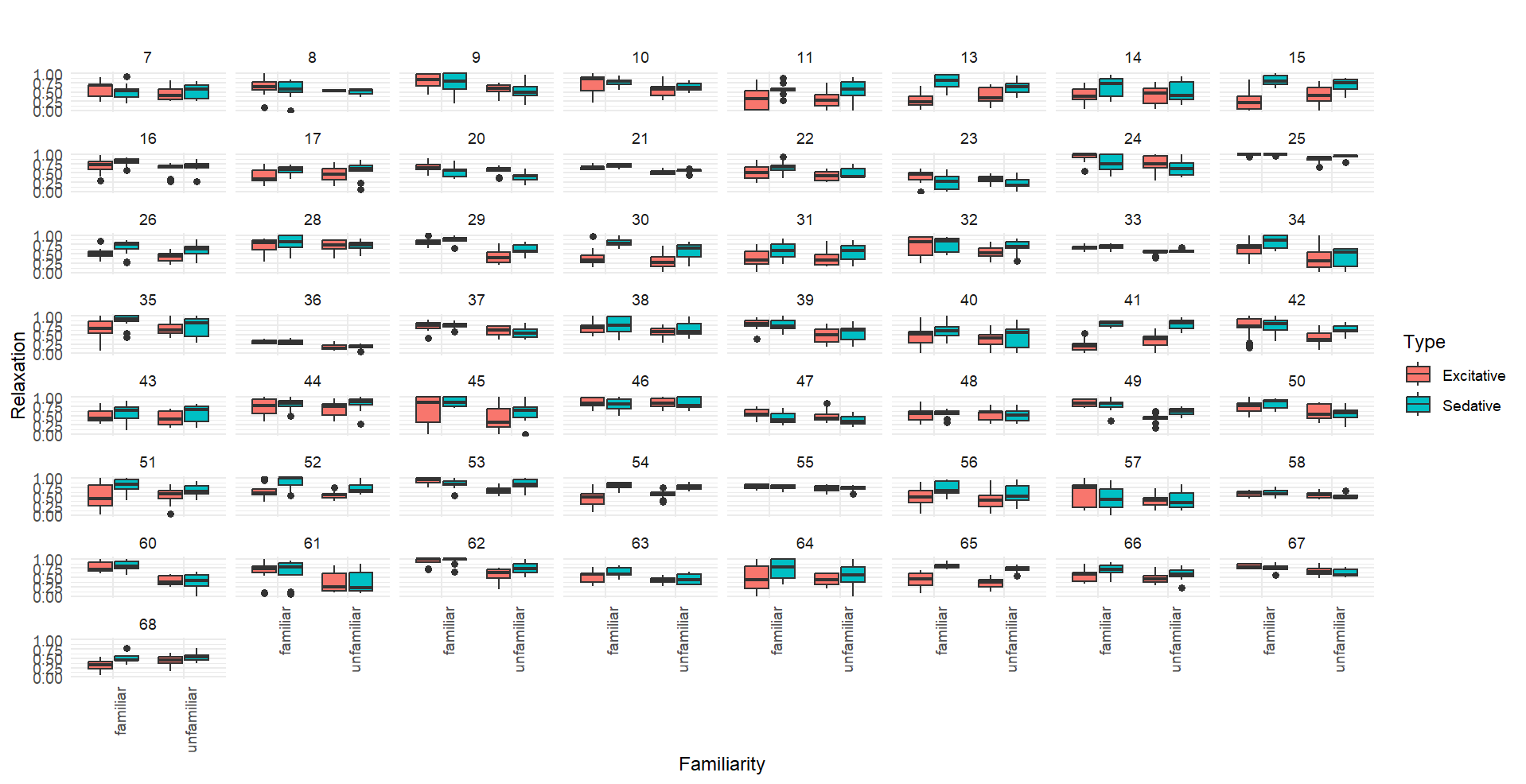

### Supplementary Figure S1: Raw relaxation responses depending on Familiarity and Type per participant.

### Clustering

| Supplementary Table S23: Participant IDs and how they split in the four clusters | |
| --- | --- |
| **Cluster** | **fMRI_id** |
| **Unclear-drive** | 7, 8, 20, 23, 36, 37, 43, 46, 47, 48, 55, 57, 58 |
| **Familiarity-driven** | 9, 10, 25, 29, 34, 39, 42, 45, 49, 50, 53, 60, 61, 62, 67 |
| **Type-driven** | 11, 13, 14, 15, 17, 31, 41, 54, 64, 65, 68 |
| **Familiarity and Type-driven** | 16, 21, 22, 26, 28, 30, 32, 33, 35, 38, 40, 44, 51, 52, 56, 63, 66 |

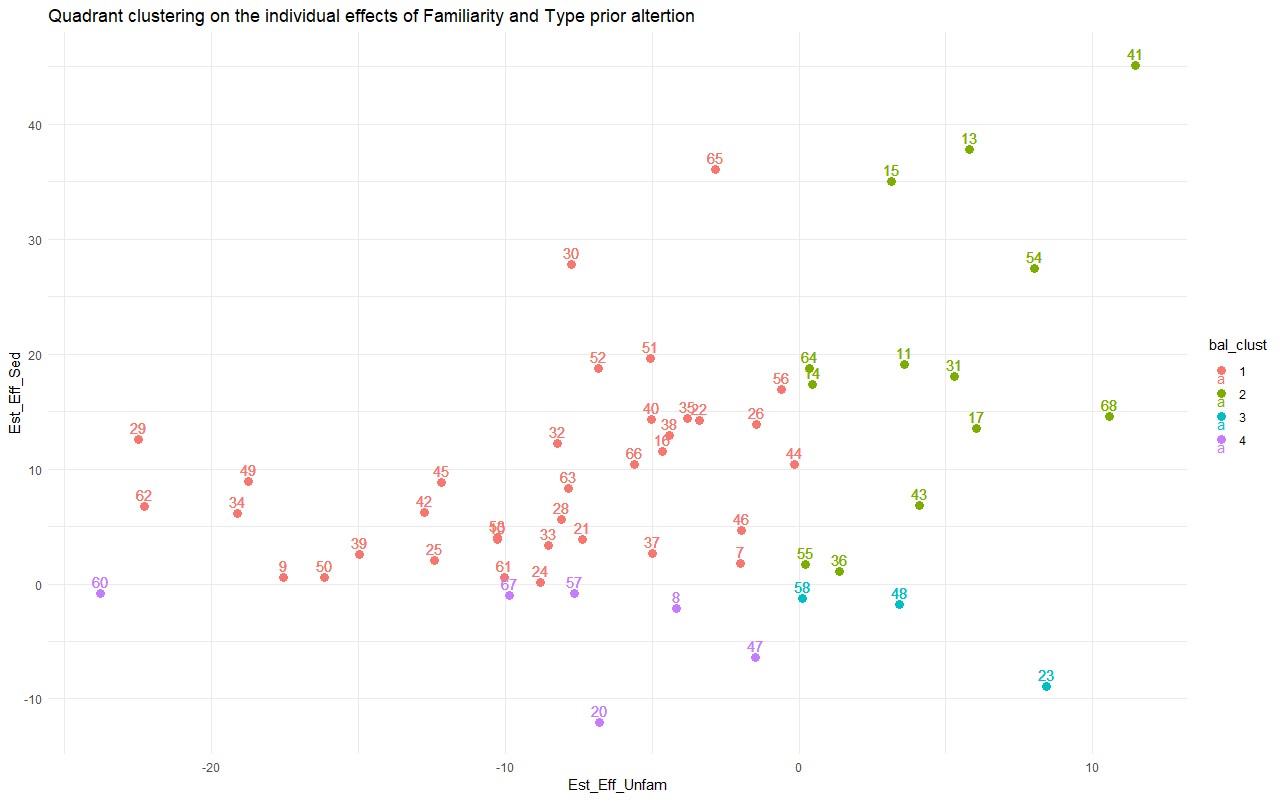

#### Supplementary Figure S2: Figure of the individual effects of Familiarity and Type with individual data points.

The colours indicate the quadrant clustering method.

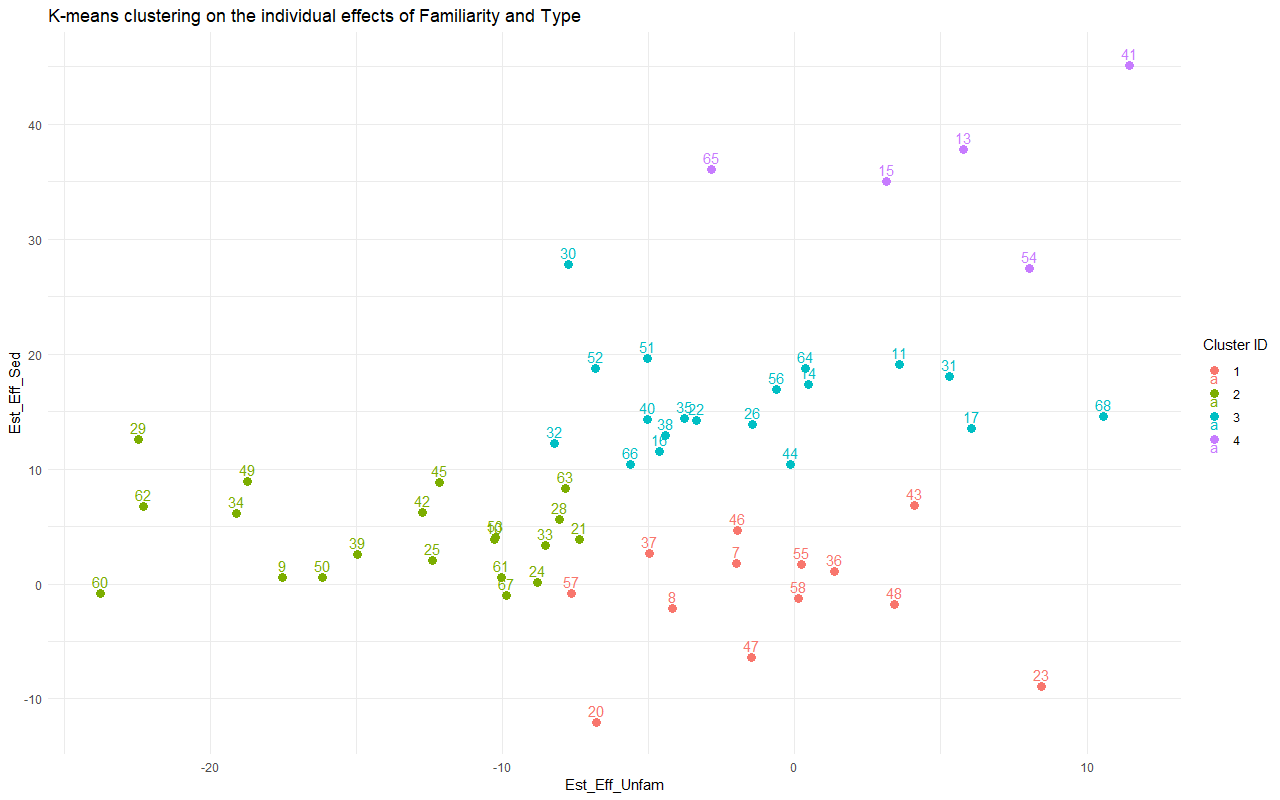

Supplementary Figure S3: Figure of the individual effects of Familiarity and Type with individual data points.
The colours indicate the k-means clustering method.

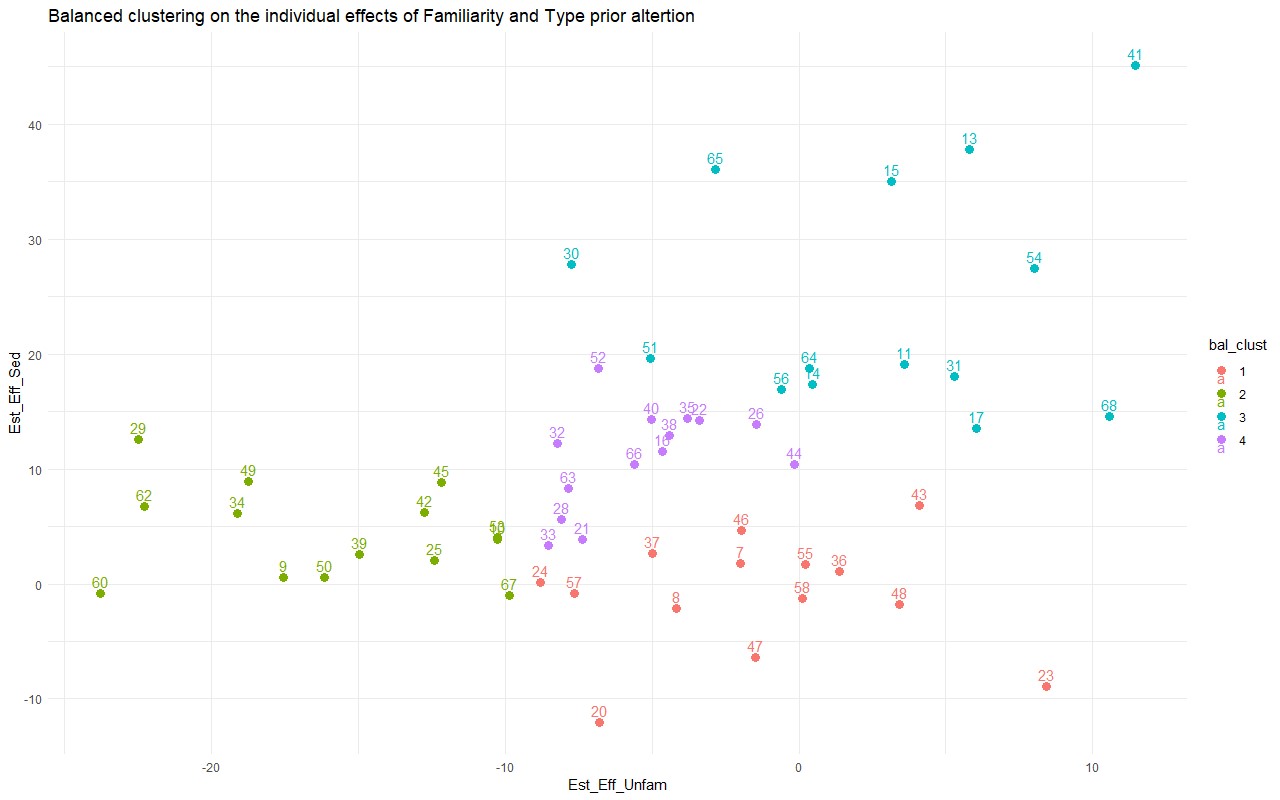

Supplementary Figure S4: Figure of the individual effects of Familiarity and Type with individual data points.
The colours indicate the balanced k-means clustering method.

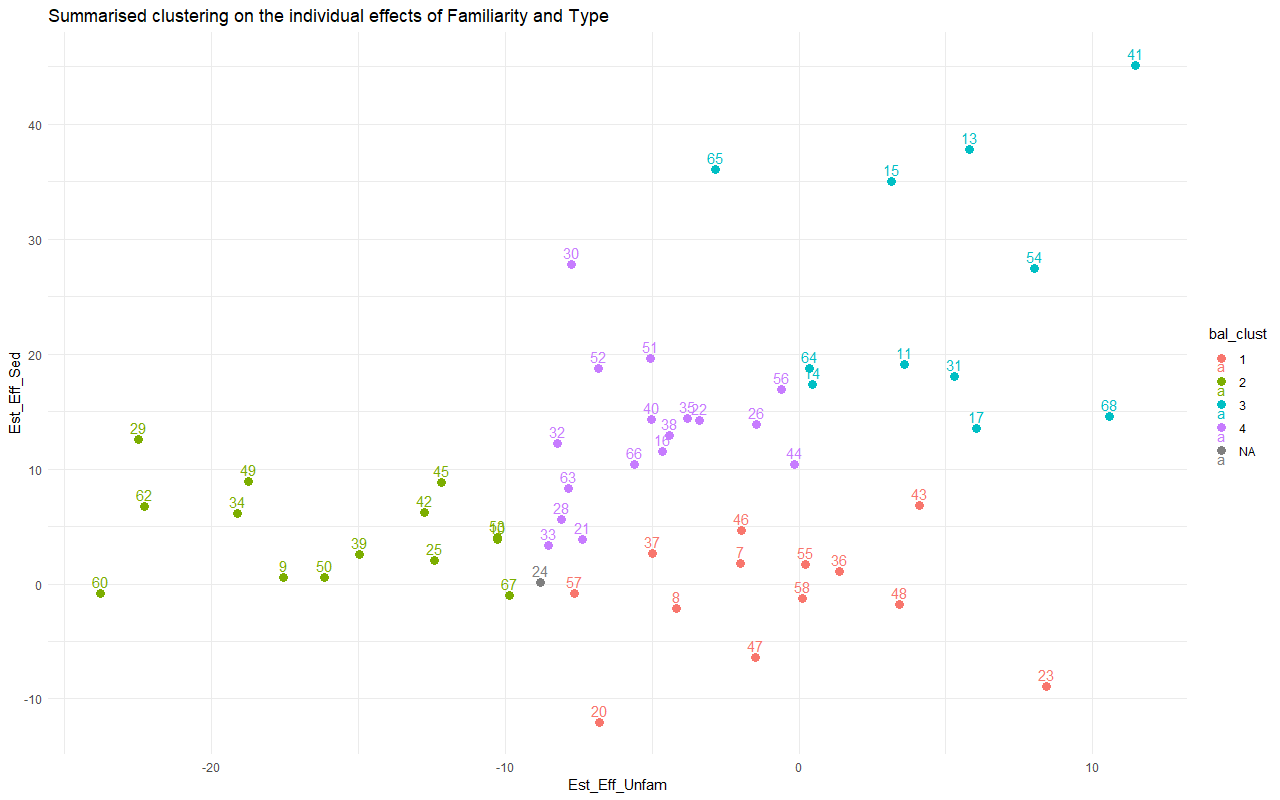

Supplementary Figure S5: Figure of the individual effects of Familiarity and Type with individual data points.
The colours indicate the summarised compound clustering based on the previous three clustering methods.

### Neuroimaging Results

| Supplementary Table S24: Brain areas included in the grey matter mask | |
| --- | --- |
| **Auditory** | Heschl’s gyrus, Superior Temporal Gyrus, Temporal Pole Superior and Middle, Middle Temporal Gyrus, Inferior Temporal Gyrus |
| **Memory** | Hippocampus, Parahippocampal gyrus, Precuneus, Angular Gryus |
| **Reward** | Amygdala, Ventral Tegmental Area, Nucleus Accumbens |
| **Motor** | Cerebellum (all), Motor Cortex (Supramarginal Gyrus, Precentral gyrus, Supplementary Motor Area), Basal Ganglia (Caudate, Putamen, Pallidum) |
| **Attention** | dorsolateral Superior frontal gyrus, Middle Frontal Gyrus, Opercular Inferior Frontal Gyrus, Triangular Inferior Frontal Gyrus, IFG pars orbitalis, Superior Parietal Gyrus, Inferior Parietal Gyrus |
| **Somatosensory** | Postcentral Gyrus, Rolandic operculum |
| **Emotion/Cognition** | Medial, Anterior, Posterior and Lateral Orbital Gyrus, Medial and Medial Orbital Superior frontal gyrus |
| **Thalamus** | All nuclei |
| **Salience Network** | Insula, Middle cingulate, Paracingulate and Post Cingulate Gyrus, Subgenual, Pregenual and Supracallosal Anterior Cingulate Cortex |
| *Areas defined according to AAL3*  *All areas are included bihemispherically* | |

| Supplementary Table S25: Main effects of Familiarity and Type for global dataset | | | | | |
| --- | --- | --- | --- | --- | --- |
| **Main Effect of Familiarity** | | | | | |
| Region | x | y | z | F | Cluster size |
| Supplementary Motor Cortex | -2 | 0 | 68 | 180.78 | 1786 |
| Supplementary Motor Cortex | -10 | 14 | 36 | 46.27 |  |
| PreCentral Gyrus | -48 | -6 | 50 | 129.32 | 748 |
| Cerebellum | 30 | -62 | -24 | 110.17 | 1043 |
| Cerebellum | 42 | -64 | -26 | 77.09 |  |
| Cerebellum | 16 | -74 | -20 | 35.88 |  |
| Temporal Pole | -50 | 12 | -14 | 91.95 | 5024 |
| Operculum | -50 | 6 | 2 | 89.8 |  |
| IFG | -52 | 32 | 2 | 73.12 |  |
| Planum Temporale | -50 | -42 | 24 | 79.29 | 556 |
| Operculum | 46 | 8 | 0 | 77.84 | 591 |
| Temporal Pole | 52 | 12 | -12 | 49.87 |  |
| PreCentral Gyrus | 52 | 0 | 48 | 68.27 | 337 |
| IFG | 54 | 32 | 2 | 57.57 | 128 |
| SFG | 24 | 22 | 60 | 54 | 301 |
| MFG | 36 | 26 | 48 | 31.04 |  |
| Cerebellum | -30 | -56 | -26 | 49.92 | 324 |
| Cerebellum | -22 | -60 | -22 | 45.56 |  |
| Cerebellum | -4 | 48 | -14 | 42.36 | 122 |
| SFG | -8 | 58 | 16 | 42.34 | 186 |
| Cingulate Gyrus | -2 | -10 | 42 | 33.71 | 36 |
| Amygdala | 18 | -4 | -12 | 31.5 | 18 |
| PreCentral Gyrus | 62 | 4 | 14 | 30.76 | 27 |
| MTG | -48 | -44 | 4 | 29.97 | 111 |
| MTG | -52 | -12 | -14 | 29.5 | 46 |
| Angular Gyrus | 54 | -52 | 40 | 29.27 | 94 |
| Putamen | 18 | 8 | 4 | 28.58 | 51 |
| SFG | -8 | 54 | 34 | 27.07 | 12 |
| STG | 48 | -34 | 4 | 25.97 | 22 |
| MFG | -24 | 38 | 22 | 25.3 | 45 |
| MFG | -30 | 44 | 28 | 24.96 |  |
| **Main Effect of Type** | | | | | |
| Region | x | y | z | F | Cluster size |
| TTG | -48 | -14 | 2 | 474.22 | 2921 |
| Insula | -28 | -22 | 8 | 24.45 |  |
| TTG | 50 | -10 | 0 | 410.33 | 2761 |
| Planum Polare | 42 | 6 | -20 | 25.28 |  |
| Cerebellum | -30 | -58 | -26 | 34.22 | 116 |
| Cerebellum | 32 | -58 | -24 | 33.69 | 191 |
| Cerebellum | 38 | -64 | -24 | 31.16 |  |
| SFG | -10 | -2 | 66 | 32.2 | 165 |
| Supplementary Motor Cortex | 6 | 0 | 72 | 26.96 |  |
| Putamen | 22 | 8 | 4 | 29.97 | 63 |
| **Main effect interaction** | | | | | |
| Region | x | y | z | F | Cluster size |
| STG | -58 | -8 | 0 | 65.09 | 639 |
| STG | 60 | -12 | 0 | 62.67 | 596 |
| STG | 56 | -30 | 4 | 31.87 |  |
| MNI, FWE p<.05, k>10, labelling with Neuromorphometrics Abbreviations: IFG/SFG/MFG=inferior/superior/middle frontal gyrus, TTG/STG= Transverse/superior temporal gyrus | | | | | |

| Supplementary Table S26: All contrasts for the global dataset | | | | | |
| --- | --- | --- | --- | --- | --- |
| **Contrast Excitative vs Sedative** | | | | | |
| Region | x | y | z | T | Cluster size |
| TTG | -48 | -14 | 2 | 21.78 | 3027 |
| Thalamus | -28 | -22 | 8 | 4.94 |  |
| TTG | 50 | -10 | 0 | 20.26 | 2859 |
| Planum Polare | 42 | 6 | -20 | 5.03 |  |
| Cerebellum | -30 | -58 | -26 | 5.85 | 146 |
| Cerebellum | 32 | -58 | -24 | 5.8 | 244 |
| Cerebellum | 38 | -64 | -24 | 5.58 |  |
| Supplementary Motor Cortex | -10 | -2 | 66 | 5.67 | 234 |
| Supplementary Motor Cortex | 6 | 0 | 72 | 5.19 |  |
| Putamen | 22 | 8 | 4 | 5.47 | 91 |
| Caudate | 20 | 20 | 2 | 5.04 |  |
| **Contrast Sedative vs Excitative** | | | | | |
| No significant peaks | | | | | |
| **Contrast Familiar vs Unfamiliar** | | | | | |
| Region | x | y | z | T | Cluster size |
| Supplementary Motor Cortex | -2 | 0 | 68 | 13.45 | 1918 |
| Cingulate Gyrus | -10 | 14 | 36 | 6.8 |  |
| PreCentral Gyrus | -48 | -6 | 50 | 11.37 | 796 |
| Cerebellum | 30 | -62 | -24 | 10.5 | 1165 |
| Cerebellum | 42 | -64 | -26 | 8.78 |  |
| Cerebellum | 16 | -74 | -20 | 5.99 |  |
| Temporal Pole | -50 | 12 | -14 | 9.59 | 5487 |
| Operculum | -50 | 6 | 2 | 9.48 |  |
| IFG | -52 | 32 | 2 | 8.55 |  |
| Planum Temporale | -50 | -42 | 24 | 8.9 | 828 |
| MTG | -48 | -44 | 4 | 5.47 |  |
| Insula | 46 | 8 | 0 | 8.82 | 676 |
| Temporal Pole | 52 | 12 | -12 | 7.06 |  |
| PreCentral Gyrus | 62 | 4 | 14 | 5.55 |  |
| PreCentral Gyrus | 52 | 0 | 48 | 8.26 | 368 |
| IFG | 54 | 32 | 2 | 7.59 | 143 |
| Cerebellum | -30 | -56 | -26 | 7.07 | 372 |
| Cerebellum | -22 | -60 | -22 | 6.75 |  |
| Cerebellum | -12 | -64 | -18 | 4.63 |  |
| Cingulate Gyrus | -4 | 48 | -14 | 6.51 | 140 |
| MFG | -8 | 58 | 16 | 6.51 | 214 |
| Cingulate Cortex | -2 | -10 | 42 | 5.81 | 44 |
| Amygdala/  Parahippocampal Gyrus | 18 | -4 | -12 | 5.61 | 24 |
| Putamen | 18 | 8 | 4 | 5.35 | 88 |
| Pallidum | 12 | 6 | -2 | 5.1 |  |
| Putamen | 24 | 6 | 14 | 4.79 |  |
| SFG | -8 | 54 | 34 | 5.2 | 18 |
| MTG | 48 | -34 | 4 | 5.1 | 42 |
| MFG | -24 | 38 | 22 | 5.03 | 74 |
| MFG | -30 | 44 | 28 | 5 |  |
| **Contrast Unfamiliar vs Familiar** | | | | | |
| Region | x | y | z | T | Cluster size |
| SFG | 24 | 22 | 60 | 7.35 | 362 |
| MFG | 36 | 26 | 48 | 5.57 |  |
| Angular Gyrus | 54 | -52 | 40 | 5.41 | 139 |
| MFG | 32 | 60 | 14 | 5.11 | 15 |
| MNI, FWE p<.05, k>10, labelling with Neuromorphometrics | | | | | |
| Abbreviations: IFG/SFG/MFG=inferior/superior/middle frontal gyrus, TTG/STG/MTG= Transverse/superior/medial temporal gyrus | | | | | |

| Supplementary Table S27: Main effects of Familiarity and Type and all contrasts for Cluster Familiarity and Type-driven | | | | | |
| --- | --- | --- | --- | --- | --- |
| **Main effect of Familiarity** | | | | | |
| Region | x | y | z | F | Cluster size |
| PreCentral Gyrus | -48 | -6 | 54 | 55.08 | 174 |
| Operculum | 44 | 8 | 2 | 47.29 | 78 |
| Supplementary Motor Cortex | -4 | -2 | 70 | 41.25 | 402 |
| Supplementary Motor Cortex | 4 | 6 | 62 | 38.62 |  |
| Operculum | -44 | 6 | 6 | 36.28 | 54 |
| PreCentral Gyrus | 48 | -2 | 48 | 31.23 | 35 |
| Cerebellum | 40 | -64 | -26 | 28.65 | 20 |
| **Main effect of Type** | | | | | |
| Region | x | y | z | F | Cluster size |
| TTG | -48 | -16 | 2 | 258.5 | 1563 |
| TTG | -36 | -28 | 12 | 74.34 |  |
| Planum Polare | -38 | -36 | 10 | 64.17 |  |
| TTG | 52 | -6 | 0 | 171.08 | 1310 |
| TTG | 46 | -14 | 8 | 142.15 |  |
| Planum Temporale | 58 | -14 | 6 | 103.19 |  |
| **Main effect of interaction** | | | | | |
| No significant peaks | | | | | |
| **Contrast Excitative vs Sedative** | | | | | |
| Region | x | y | z | T | Cluster size |
| TTG | -48 | -16 | 2 | 16.08 | 1656 |
| TTG | -36 | -28 | 12 | 8.62 |  |
| Planum Temporale | -38 | -36 | 10 | 8.01 |  |
| Planum Polare | 52 | -6 | 0 | 13.08 | 1403 |
| TTG | 46 | -14 | 8 | 11.92 |  |
| Planum Temporale | 58 | -14 | 6 | 10.16 |  |
| **Contrast Sedative vs Excitative** | | | | | |
| No significant peaks | | | | | |
| **Contrast Familiar vs Unfamiliar** | | | | | |
| Region | x | y | zmm | T | equivk |
| PreCentral Gyrus | -48 | -6 | 54 | 7.42 | 206 |
| Operculum | 44 | 8 | 2 | 6.88 | 100 |
| Supplementary Motor Cortex | -4 | -2 | 70 | 6.42 | 485 |
| Supplementary Motor Cortex | 4 | 6 | 62 | 6.21 |  |
| Operculum | -44 | 6 | 6 | 6.02 | 78 |
| PreCentral Gyrus | 48 | -2 | 48 | 5.59 | 50 |
| Cerebellum | 40 | -64 | -26 | 5.35 | 57 |
| Cerebellum | 46 | -54 | -32 | 5.06 |  |
| Cerebellum | 26 | -60 | -24 | 5.34 | 15 |
| **Contrast Unfamiliar vs Familiar** | | | | | |
| No significant peaks | | | | | |
| MNI, FWE p<.05, k>10, labelling with Neuromorphometrics Abbreviations: TTG= Transverse temporal gyrus | | | | | |

| Supplementary Table S28: Main effects of Familiarity and Type and all contrasts for Cluster Familiarity-driven | | | | | |
| --- | --- | --- | --- | --- | --- |
| **Main effect of Familiarity** | | | | | |
| Region | x | y | z | F | Cluster size |
| Supplementary Motor Cortex | -2 | 0 | 66 | 77.26 | 440 |
| PreCentral Gyrus | -50 | -6 | 48 | 45.29 | 76 |
| Cerebellum | 26 | -62 | -22 | 37.24 | 76 |
| Cerebellum | 34 | -60 | -24 | 34.63 |  |
| Operculum | -48 | 4 | 4 | 34.53 | 32 |
| MFG | 42 | 42 | 24 | 31.39 | 24 |
| Thalamus | -16 | -10 | 16 | 31.1 | 11 |
| **Main effect of Type** | | | | | |
| Region | x | y | z | F | Cluster size |
| TTG | -46 | -14 | 2 | 71.94 | 460 |
| Planum Polare | 52 | -4 | -2 | 65.12 | 549 |
| TTG | 42 | -16 | 4 | 64.27 |  |
| TTG | 42 | -24 | 10 | 39.6 |  |
| **Main effect of interaction** | | | | | |
| No significant peaks | | | | | |
| **Contrast Excitative vs Sedative** | | | | | |
| Region | x | y | z | T | Cluster size |
| TTG | -46 | -14 | 2 | 8.48 | 538 |
| Planum Polare | 52 | -4 | -2 | 8.07 | 639 |
| TTG | 42 | -16 | 4 | 8.02 |  |
| TTG | 42 | -24 | 10 | 6.29 |  |
| **Contrast Sedative vs Excitative** | | | | | |
| No significant peaks | | | | | |
| **Contrast Familiar vs Unfamiliar** | | | | | |
| Region | x | y | z | T | Cluster size |
| Supplementary Motor Cortex | -2 | 0 | 66 | 8.79 | 503 |
| PreCentral Gyrus | -50 | -6 | 48 | 6.73 | 99 |
| Cerebellum | 26 | -62 | -22 | 6.1 | 115 |
| Cerebellum | 34 | -60 | -24 | 5.88 |  |
| Operculum | -48 | 4 | 4 | 5.88 | 54 |
| MFG | -16 | -10 | 16 | 5.58 | 24 |
| Thalamus | -42 | 28 | -4 | 5.14 | 12 |
| **Contrast Unfamiliar vs Familiar** | | | | | |
| Region | x | y | z | T | Cluster size |
| MFG | 42 | 42 | 24 | 5.6 | 34 |
| MNI, FWE p<.05, k>10, labelling with Neuromorphometrics Abbreviations: MFG= middle frontal gyrus, TTG = Transverse temporal gyrus | | | | | |

| Supplementary Table S29: Main effects of Familiarity and Type and all contrasts for Cluster Type-driven | | | | | |
| --- | --- | --- | --- | --- | --- |
| **Main effect of Familiarity** | | | | | |
| Region | x | y | z | F | Cluster size |
| Operculum | -40 | 28 | 4 | 38.05 | 35 |
| **Main effect of Type** | | | | | |
| Region | x | y | z | F | Cluster size |
| TTG | -46 | -16 | 4 | 74.26 | 604 |
| Planum Polare | -52 | -2 | 0 | 63.33 |  |
| Planum Temporale | -50 | -30 | 10 | 34.68 |  |
| TTG | 48 | -14 | 4 | 68.12 | 439 |
| Planum Polare | 54 | 0 | -4 | 39.42 |  |
| Planum Temporale | 58 | -22 | 8 | 32.76 |  |
| **Main effect of interaction** | | | | | |
| No significant peaks | | | | | |
| **Contrast Excitative vs Sedative** | | | | | |
| Region | x | y | z | T | Cluster size |
| TTG | -46 | -16 | 4 | 8.62 | 723 |
| Planum Polare | -52 | -2 | 0 | 7.96 |  |
| Planum Temporale | -50 | -30 | 10 | 5.89 |  |
| TTG | 48 | -14 | 4 | 8.25 | 541 |
| Planum Polare | 54 | 0 | -4 | 6.28 |  |
| Planum Temporale | 58 | -22 | 8 | 5.72 |  |
| **Contrast Sedative vs Excitative** | | | | | |
| No significant peaks | | | | | |
| **Contrast Familiar vs Unfamiliar** | | | | | |
| Region | x | y | z | T | Cluster size |
| Operculum | -40 | 28 | 4 | 6.17 | 55 |
| **Contrast Sedative vs Excitative** | | | | | |
| No significant peaks | | | | | |
| MNI, FWE p<.05, k>10, labelling with Neuromorphometrics Abbreviations: TTG= Transverse temporal gyrus | | | | | |

| Supplementary Table S30: Main effects of Familiarity and Type and all contrasts for Cluster Unclear-drive | | | | | |
| --- | --- | --- | --- | --- | --- |
| **Main effect of Familiarity** | | | | | |
| Region | x | y | z | F | Cluster size |
| Supplementary Motor Cortex | 0 | 0 | 68 | 55.44 | 197 |
| PreCentral Gyrus | -48 | 0 | 52 | 46.8 | 39 |
| Temporal Pole | -46 | 10 | -18 | 45.73 | 126 |
| Temporal Pole | -54 | 12 | -4 | 38.78 |  |
| STG | -56 | 4 | -10 | 30.7 |  |
| PreCentral Gyrus | 40 | 0 | 44 | 36.04 | 46 |
| PreCentral Gyrus | 50 | 4 | 50 | 34.74 |  |
| Supramarginal Gyrus | -52 | -44 | 26 | 29.99 | 14 |
| **Main effect of Type** |  |  |  |  |  |
| Region | x | y | z | F | Cluster size |
| TTG | 50 | -8 | 2 | 202.51 | 1069 |
| STG | 48 | -30 | 8 | 35.26 |  |
| STG | 58 | -24 | 6 | 33.72 |  |
| Planum Polare | -50 | -12 | 2 | 120.18 | 1256 |
| Planum Temporale | -54 | -22 | 6 | 83.79 |  |
| **Main effect of interaction** | | | | | |
| No significant peaks | | | | | |
| **Contrast Excitative vs Sedative** | | | | | |
| Region | x | y | z | T | Cluster size |
| TTG | 50 | -8 | 2 | 14.23 | 1196 |
| STG | 48 | -30 | 8 | 5.94 |  |
| STG | 58 | -24 | 6 | 5.81 |  |
| Planum Polare | -50 | -12 | 2 | 10.96 | 1352 |
| Planum Temporale | -54 | -22 | 6 | 9.15 |  |
| **Contrast Sedative vs Excitative** | | | | | |
| No significant peaks | | | | | |
| **Contrast Familiar vs Unfamiliar** | | | | | |
| Region | x | y | z | T | Cluster size |
| Supplementary Motor Cortex | 0 | 0 | 68 | 7.45 | 235 |
| PreCentral Gyrus | -48 | 0 | 52 | 6.84 | 62 |
| Temporal Pole | -46 | 10 | -18 | 6.76 | 168 |
| Temporal Pole | -54 | 12 | -4 | 6.23 |  |
| STG | -56 | 4 | -10 | 5.54 |  |
| PreCentral Gyrus | 40 | 0 | 44 | 6 | 93 |
| PreCentral Gyrus | 50 | 4 | 50 | 5.89 |  |
| Temporal Pole | 52 | 12 | -12 | 5.59 | 21 |
| Cerebellum | 34 | -62 | -24 | 5.51 | 26 |
| STG | -62 | -50 | 12 | 5.5 | 24 |
| Supramarginal Gyrus | -52 | -44 | 26 | 5.48 | 36 |
| **Contrast Sedative vs Excitative** | | | | | |
| No significant peaks | | | | | |
| MNI, FWE p<.05, k>10, labelling with Neuromorphometrics Abbreviations: STG/MTG= superior/medial temporal gyrus | | | | | |

| Supplementary Table S31: Descriptive demographic information per cluster | | | | |
| --- | --- | --- | --- | --- |
|  | Familiarity and Type driven (N= 17) | Familiarity-driven (N= 15) | Type-driven  (N= 11) | Unclear-drive (N=13) |
| Nationality | Danish: 4  Western Europe: 6  Eastern Europe: 4  Non-Western: 3 | Danish: 2  Western Europe: 7  Eastern Europe: 6  Non-Western: 0 | Danish: 3  Western Europe: 3  Eastern Europe: 3  Non-Western: 2 | Danish: 2  Western Europe: 3  Eastern Europe: 5  Non-Western: 3 |
| Mean age (SD) | 24.94(4.40) | 23.6(3.24) | 27.18(4.47) | 27.52(5.44) |
| Gender | Female: 12  Male: 5  Non-binary: 0 | Female: 11  Male: 4  Non-binary: 0 | Female: 6  Male: 4  Non-binary: 1 | Female: 9  Male: 4  Non-binary:0 |
| Handedness | Left: 2  Right: 14  Ambidextrous: 1 | Left: 0  Right: 14  Ambidextrous: 1 | Left: 0  Right: 11  Ambidextrous: 0 | Left: 0  Right: 11  Ambidextrous: 2 |
| Education | High School: 5  Bachelor: 7  Higher: 5 | High School: 6  Bachelor: 4  Higher: 5 | High School: 4  Bachelor: 4  Higher: 3 | High School: 2  Bachelor: 6  Higher: 5 |
| Singing ability | Poor: 3  Medium: 6  Ok in group: 5  Good: 3 | Poor: 5  Medium: 5  Ok in group: 3  Good: 2 | Poor: 0  Medium: 4  Ok in group: 4  Good: 3 | Poor: 2  Medium: 4  Ok in group: 4  Good: 3 |
| Frequency music relax | Rarely: 1  Sometimes:3  Often: 6  Always: 7 | Rarely: 0  Sometimes: 2  Often: 7  Always: 6 | Rarely: 0  Sometimes: 2  Often: 4  Always: 5 | Rarely: 0  Sometimes: 2  Often: 8  Always: 3 |
| Mean extraversion (SD) | 5.82(2.69) | 6(3.31) | 5.64(2.77) | 6.64(2.86) |
| Mean Music Training | 2.73(1.65) | 2.33(1.82) | 2.67(1.26) | 1.95(1.28) |
| Music genres | Country: 1  EDM: 2  Folk: 3  Funk: 1  House: 1  Indie: 11  Jazz: 4  Lo.Fi: 7  Meditation: 0  New.age: 1  Pop: 16  Punk: 0  R.B: 8  Rap: 4  Reggae:1  Rock: 9  Soundtrack: 3  World: 1 | Country: 1  EDM: 2  Folk: 0  Funk: 2  House: 3  Indie: 11  Jazz: 5  Lo.Fi: 3  Meditation: 2  New.age: 1  Pop: 9  Punk: 1  R.B: 5  Rap: 7  Reggae: 0  Rock: 9  Soundtrack: 3  World: 3 | Country: 1  EDM: 2  Folk: 5  Funk: 3  House: 2  Indie: 8  Jazz: 2  Lo.Fi: 1  Meditation: 3  New age:  Pop: 8  Punk: 1  R.B: 4  Rap: 5  Reggae: 2  Rock: 7  Soundtrack: 5  World: 1 | Country: 3  EDM: 4  Folk: 6  Funk: 0  House: 4  Indie: 3  Jazz: 7  Lo.Fi: 5  Meditation: 3  New age: 4  Pop: 12  Punk: 2  R&B: 2  Rap:0  Reggae: 2  Rock: 9  Soundtrack: 5  World: 3 |
| Type music relax | Neo-soul: 1  Lofi: 6  Indie: 8  Psychedelic: 2  Instrumental: 4  Acoustic: 3  Alternative Rock: 1  Binaural Beats: 1  Classic Rock: 2  Opera: 1  Classical Music: 3  Meditation Music: 2  R&B: 1  Pop: 6  Country: 2  Jazz: 4  Chillwave: 1  Electronic Music: 3  Downtempo: 1  Folk: 3  IDM: 1  Rock: 4  Blues: 1  Sad or Slow Music: 3  Soft Pop: 1  Soft Rock: 1  Soul: 1  Soundtrack: 2  Trance: 1  Metal: 1  Oriental: 1  Kpop: 1  Jpop: 1  Rap: 1 | 8nos and 9nos music: 1  Acoustic: 2  Alternative Rock: 1  Binaural Beats: 1  Chillwave: 1  Classical: 4  Country: 1  Downtempo: 1  Electronic: 2  Folk: 1  Hungarian Folk Pop: 1  Indie: 3  Instrumental: 5  Jazz: 3  Kpop: 1  Jpop: 1  Lo-Fi: 4  Meditation: 2  Metal: 1  Nostalgic Music: 1  Opera: 1  Oriental: 1  Piano Music: 1  Pop: 5  Psychedelic: 2  Rap: 2  Rock: 3  Soft Pop: 1  Soundtrack: 1  Slow Pop: 1  Trance: 1  Weirdcore: 1 | Acoustic: 1  Classical: 4  Country: 2  Electronic: 2  Folk: 1  Indie: 1  Instrumental: 4  Jazz: 2  Jpop: 1  Kpop: 1  Lo-Fi: 3  Meditation: 1  Metal: 1  Opera: 1  Oriental: 1  Pop: 1  Psychedelic: 1  Rap: 1  Rock: 3  Slow: 2  Soundtrack: 1  Trance: 1 | Acoustic: 1  Classical: 3  Country: 2  Electronic: 2  Folk: 2  IDM: 1  Indie: 5  Instrumental: 5  Jazz: 5  Lofi: 4  Meditation: 2  Opera: 1  Piano: 1  Pop: 4  Psychedelic: 2  Rap: 1  Rock: 6  Slow: 2  Soul: 1  Soundtrack: 2  Trance: 1 |

| Supplementary Table S32: Selected Stimuli | | | | |
| --- | --- | --- | --- | --- |
| Type | Familiarity | Track Artist and Title | Rated Familiarity Mean(SD) | Spotify Energy |
| Excitative | Familiar | Whitney Houston - I Wanna Dance With Somebody | 98.5(5.88) | 0.82 |
|  |  | Hozier - Take Me To Church | 98.35(5.84) | 0.66 |
|  |  | Bon Jovi - Living On A Prayer | 97.4(8.30) | 0.89 |
|  |  | Coldplay - A Sky Full Of Stars | 96.85(6.90) | 0.68 |
|  |  | Spice Girls - Wannabe | 96.15(19.61) | 0.86 |
|  |  | Abba - Dancing Queen | 95.77(19.63) | 0.87 |
|  |  | Alicia Keys - Girl on Fire | 93.69(14.70) | 0.70 |
|  |  | The Fray - How to Save a Life | 93.58(13.06) | 0.74 |
|  |  | Owl City - Fireflies | 92.12(24.59) | 0.66 |
|  |  | Katrina & The Waves - Walking On Sunshine | 92.07(21.37) | 0.87 |
|  |  | Journey - Don't Stop Believin' | 91.69(20.54) | 0.75 |
|  |  | Maroon 5 - Payphone | 91.19(21.49) | 0.75 |
|  |  | Bruno Mars - Just the Way You Are | 91.04(26.43) | 0.84 |
|  |  | OneRepublic - Counting Stars | 90.88(20.51) | 0.71 |
|  |  | a-ha - Take On Me | 89.19(27.84) | 0.90 |
|  |  | R.E.M. - Losing My Religion | 86.88(28.96) | 0.86 |
|  |  | Elton John, Kiki Dee - Don't Go Breaking My Heart | 87.73(28.00) | 0.84 |
|  |  | Lana Del Rey - Summertime Sadness | 84.19(29.96) | 0.65 |
|  | Unfamiliar | Nona - Givin It All | 21.77(27.11) | 0.59 |
|  |  | David Blair - Did You Know | 12.96(18.88) | 0.72 |
|  |  | Chris Christian - Walnut Hill | 17.54(28.48) | 0.57 |
|  |  | The goo goo dolls - Bringing on the Light | 14.42(22.90) | 0.88 |
|  |  | Jasmine Thompson - Do It Now | 24.62(28.81) | 0.78 |
|  |  | The pointer sisters - What a Surprise | 10.42(16.22) | 0.72 |
|  |  | Stefanie Heinzmann - Revolution | 14.42(20.20) | 0.58 |
|  |  | You Me At Six - No One Does It Better | 18.35(26.68) | 0.62 |
|  |  | Badly Drawn Boy - The Time Of Times | 13.69(19.64) | 0.63 |
|  |  | Lucy Spraggan - Run | 23.19(29.31) | 0.61 |
|  |  | Brian Elliot - Room to Grow | 15.35(24.72) | 0.63 |
|  |  | Spawnbreezie - All Alone | 11.15(20.46) | 0.81 |
|  |  | Kawala - Arms Wide Open | 16.54(21.25) | 0.74 |
|  |  | Lovebugs - The Aftermoon | 16.19(23.26) | 0.64 |
|  |  | The Kooks - Always Free | 16.35(23.63) | 0.85 |
|  |  | The Wombats - Last Night I Dreamt... | 15.65(25.34) | 0.81 |
|  |  | Simply Red - Thinking of You | 24.46(31.56) | 0.59 |
|  |  | Lorren - Love Definition | 7.81(11.93) | 0.70 |
| Sedative | Familiar | Celine Dion - My Heart Will Go On | 99.54(1.55) | 0.28 |
|  |  | Ben E. King - Stand By Me | 98.31(4.45) | 0.23 |
|  |  | John Legend - All of Me | 92.88(19.95) | 0.26 |
|  |  | John Lennon - Imagine | 91.88(22.63) | 0.26 |
|  |  | Bobby McFerrin - Don't Worry Be Happy | 86.88(24.26) | 0.17 |
|  |  | Phil Collins - In The Air Tonight | 86.27(27.90) | 0.24 |
|  |  | A Great Big World, Christina Aguilera - Say Something | 84.65(34.59) | 0.15 |
|  |  | James Bay - Let It Go | 81.38(31.76) | 0.21 |
|  |  | Cyndi Lauper - True Colors | 76.69(34.94) | 0.21 |
|  |  | Calum Scott - Dancing On My Own | 74.27(37.48) | 0.17 |
|  |  | gnash - i hate u, i love u ft. olivia o'brien | 73.15(31.13) | 0.28 |
|  |  | Billie Eilish - No Time To Die | 72.35(39.22) | 0.22 |
|  |  | Ed Sheeran - I See Fire | 70.96(40.33) | 0.055 |
|  |  | Sam Smith - Lay Me Down | 70.46(39.73) | 0.19 |
|  |  | Simon & Garfunkel - The Sounds of Silence | 70.16(39.82) | 0.22 |
|  |  | Beatles - Yesterday Remastered 2009 | 69.62(40.29) | 0.19 |
|  |  | Whitney Houston - Saving All My Love For You | 64.58(32.94) | 0.26 |
|  |  | Air Supply - All Out Of Love | 64.04(35.31) | 0.26 |
|  | Unfamiliar | Rita Coolidge - Cherokee | 15.77(21.35) | 0.27 |
|  |  | John Hiatt - Icy Blue Heart | 13.04(21.96) | 0.16 |
|  |  | Doe Paoro - Outlines | 15.04(23.48) | 0.18 |
|  |  | Joshua Radin - Falling | 15.96(23.43) | 0.25 |
|  |  | Glenn Frey - Some Kind Of Blue | 16.85(24.68) | 0.27 |
|  |  | Crosby, Stills and Nash - House of Broken Dreams | 9.42(17.79) | 0.22 |
|  |  | Hannah Cohen - Don't Say | 11.46(20.52) | 0.19 |
|  |  | Newton Faulkner - Here Tonight | 15.69(26.38) | 0.19 |
|  |  | Rita Coolidge - Love Lessons | 9.58(15.37) | 0.26 |
|  |  | Judie Tzuke - Hurt in Your Heart | 11.5(16.23) | 0.22 |
|  |  | Priscilla Ahn - In My Bed | 13.66(23.45) | 0.26 |
|  |  | Celine Cairo - Hibernate | 11.77(19.57) | 0.24 |
|  |  | Black Pool - Paperships | 14.96(18.09) | 0.19 |
|  |  | The amazing rhythm aces - Red to Blue (When Dreams Come True) | 13.46(19.80) | 0.25 |
|  |  | Joe Crocker - Every Time it Rains | 14.35(20.02) | 0.27 |
|  |  | The war within - The Stand | 23.69(31.21) | 0.16 |
|  |  | JC Lodge - Stealing Love (Bonus Track) | 19.19(25.88) | 0.082 |
|  |  | Michael Barr - 2 Pills | 19.81(28.78) | 0.25 |
